# Mcadet: a feature selection method for fine-resolution single-cell RNA-seq data based on multiple correspondence analysis and community detection

**DOI:** 10.1101/2023.07.26.550732

**Authors:** Saishi Cui, Sina Nassiri, Issa Zakeri

## Abstract

Single-cell RNA sequencing (scRNA-seq) data analysis faces numerous challenges, including high sparsity, a high-dimensional feature space, technical biases, and biological noise. These challenges hinder downstream analysis, necessitating the use of feature selection methods to address technical biases, identify informative genes, and reduce data dimensionality. However, existing methods for selecting highly variable genes (HVGs) exhibit limited overlap and inconsistent clustering performance across benchmark datasets. Moreover, these methods often struggle to accurately select HVGs from fine-resolution scRNA-seq datasets and rare cell types, raising concerns about the reliability of their results. To overcome these limitations, we propose a novel feature selection framework for unique molecular identifiers (UMIs) scRNA-seq data called Mcadet. Mcadet integrates Multiple Correspondence Analysis (MCA), graph-based community detection, and a novel statistical testing approach. To assess the effectiveness of Mcadet, we conducted extensive evaluations using both simulated and real-world data, employing unbiased metrics for comparison. Our results demonstrate the superior performance of Mcadet in the selection of HVGs in scenarios involving fine-resolution scRNA-seq datasets and datasets containing rare cell populations. By addressing the challenges of feature selection in scRNA-seq analysis, Mcadet provides a valuable tool for improving the reliability and accuracy of downstream analyses in single-cell transcriptomics.

## 1. Introduction

Single-cell RNA sequencing (scRNA-seq) has emerged as a powerful tool for characterizing complex human or animal tissues and cell types, enabling the examination of RNA expression differences at a single-cell resolution. Recent advancements in scRNA-seq techniques have significantly contributed to our understanding of biological systems by allowing simultaneous measurement of transcript levels in thousands of individual cells [1, 2]. This approach overcomes the limitations of bulk RNA sequencing, which lacks individual resolution.

As of today, scRNA-seq experiments can be classified under two major paradigms. The most common approaches performed on the fashionable 10X Chromium [3] platform use microscopic droplets or wells to segregate an extensive number of cells, followed by shallow sequencing of the libraries [4, 5]. One of the key advantages of this technology is its capability to support unique molecular identifiers (UMIs), which are short barcodes attached to transcripts before amplification to enable more accurate estimates of expression levels and the elimination of duplicate polymerase chain reactions [6]. However, there is one major limitation of the platforms, which is the fact that each messenger RNA (mRNA) can only be sequenced at the 5’ or 3’ end. Alternatively, some other studies use the approach of concentrating relatively few cells and then sequencing them at much greater depth. These low-throughput, high-depth experiments typically isolate cells into individual wells and apply the Smart-seq2 protocol [7].

A crucial component of scRNA-seq analysis is the expression matrix, which represents the number of transcripts detected for each gene and cell. The analysis workflow of scRNA-seq data can be divided into two main sections: pre-processing and downstream analysis. Pre-processing involves quality control, normalization, data correction, feature selection, and visualization, while downstream analysis includes clustering, differential expression analysis, annotation, and gene dynamics analysis [8, 9]. However, scRNA-seq data face challenges such as high dropout rates, noise, and technical variabilities, which can impede downstream clustering and classification analysis [10]. Additionally, scRNA-seq datasets often contain a large number of genes, many of which may not provide relevant information for a specific analysis. Therefore, feature selection plays a vital role in working with multidimensional scRNA-seq datasets.

Numerous methods have been proposed for the selection of highly variable genes (HVGs) in scRNA-seq data. However, these methods often exhibit limited agreement in the genes identified as highly variable and demonstrate inconsistent clustering performance when applied to different datasets. Currently, there is no consensus on which method outperforms the others. In a study conducted by Yip et al. [11], a comparison of seven feature selection (FS) methods for scRNA-seq data revealed substantial differences among the approaches, with each tool demonstrating optimal performance under specific scenarios. This highlights the need to carefully consider the choice of FS method based on the specific characteristics and objectives of the dataset being analyzed. Furthermore, existing methods are limited to fixed resolutions and lack the ability to handle fine-resolution datasets.

In this paper, we introduce Mcadet with UMI counts as a novel feature selection framework inspired by Cell-ID [12] and corral [13]. Cell-ID is a gene signature extraction method that enhances the biological interpretation at the individual cell level, enabling the identification of novel cell types or states. On the other hand, corral is a dimensionality reduction technique specifically designed for scRNA-seq data, demonstrating superior clustering performance compared to the standard corresponding analysis (CA) and glmPCA approaches. Both of the methods rely on CA or multiple corresponding analysis (MCA), which is a statistical technique utilizing singular-value decomposition (SVD). This technique enables the simultaneous projection of individuals and variables into a shared low-dimensional space, making it suitable for non-negative, count-based data to investigate the relationship between samples and categorical variables. CA has a rich historical background, and numerous variations and extensions have been developed for diverse research contexts across various fields [14, 15]. However, its utilization in scRNA-seq data analysis has been relatively limited until recently when the two aforementioned papers introduced its application. Therefore, it is highly valuable to explore the potential of applying CA in this particular field.

Mcadet utilizes Leiden community detection and MCA to select informative genes from scRNA-seq data and facilitate cell population recovery. Our framework aims to accurately select informative genes, handle rare cell populations and fine-resolution datasets, and allow parameter customization for various resolutions, thus addressing the limitations of current approaches. **Figure 1** summarizes our feature selection framework. In general, our method encompasses five major steps for analysis: (1). Matrix pre-processing. (2). MCA decomposition. (3). Community detection for cell clustering. (4). Computation and ranking of Euclidean distances. (5). Statistical testing. Workflow details and computational details are described in the Section 4 and **Supplementary Methods**.

**Figure 1.**
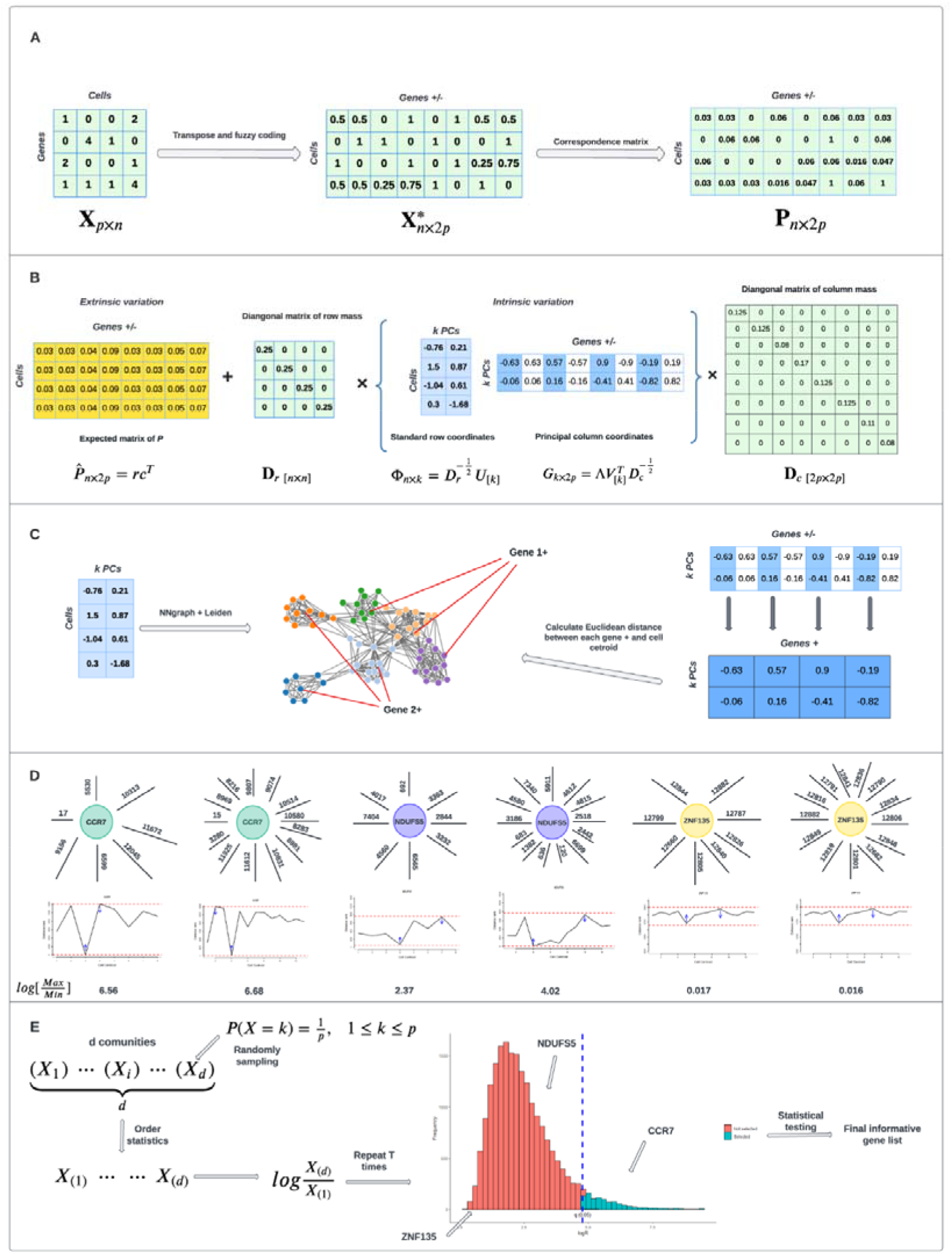
Schematic of Mcadet workflow. A-D are the five steps of Mcadet, matrix pre-processing; MCA decomposition; community detection (cells clustering); calculating and ranking the Euclidean distances and statistical testing. D gives three examples of gene distances rank pattern, for each gene, we give two examples of distance rank pattern from coarse to fine resolution. “CCR7” is a well-known driver gene for Naive CD4+ T cells. “NDUFS5” gene is a housekeeping gene, which is highly expressed in many cell types. “ZNF135” is an irrelevant gene which is only expressed in a little of cells. The statistic of log (Max rank / Min rank) in is used for selecting genes.

In order to evaluate its performance, we conducted a comparative analysis of Mcadet against seven established feature selection methods for scRNA-seq data, as outlined in Section 2. Our assessment involved applying these feature selection methods to a variety of simulated and real-world datasets. To ensure unbiased evaluation, we employed several objective metrics, such as the Jaccard similarity index for accuracy, silhouette score, purity, adjusted Rand index, and other measures for assessing clustering performance. Our results demonstrate that Mcadet outperforms other methods, particularly in fine-resolution or difficult-to-differentiate scRNA-seq datasets, as well as in datasets with rare cell populations. Overall, our novel feature selection algorithm, Mcadet, offers an improved toolbox for scRNA-seq analysis, providing more accurate feature selection and enhanced capabilities for handling challenging scRNA-seq datasets.

In Section 2, we begin with present an in-depth review of existing feature selection approaches for scRNA-seq analysis. In Section 3, we detail the datasets employed, encompassing both artificial and real-world datasets. The detailed workflow of Mcadet, the approach to conducting the feature selection method comparison, and the description of evaluation metrics are presented in Section 4. The results of the feature selection performance comparison on both artificial and real-world datasets are displayed in Section 5. Finally, in Section 6, we conclude the study with a discussion on the proposed feature selection framework, highlighting its advantages and limitation.

## 2. Overview of Feature Selection Methods

We provide a comprehensive review of feature selection approaches used in existing tools and software for scRNA-seq analysis, as summarized in **Table 1**. These approaches can be categorized into three main types: dispersion-based, dropout-rate-based, and machine-learning-based. Initially, dispersion-based methods were proposed and incorporated into scRNA-seq analysis tools such as Brennecke [16], scran [17], scVEGs [18], Seurat [19], and Scry [20]. The method introduced by Brennecke et al. [16] involves fitting a generalized linear model to a plot of the mean and squared coefficient of variation (CV2). This model enables the derivation of a coefficient of biological variation, and a chi-square test is then used to identify genes with high variance in their coefficient of biological variation, indicating potential variability in gene expression. Subsequent modifications led to the development of different feature selection methods, such as those implemented in scran, scVEGs, Seurat, and Scry.

**Table 1.**
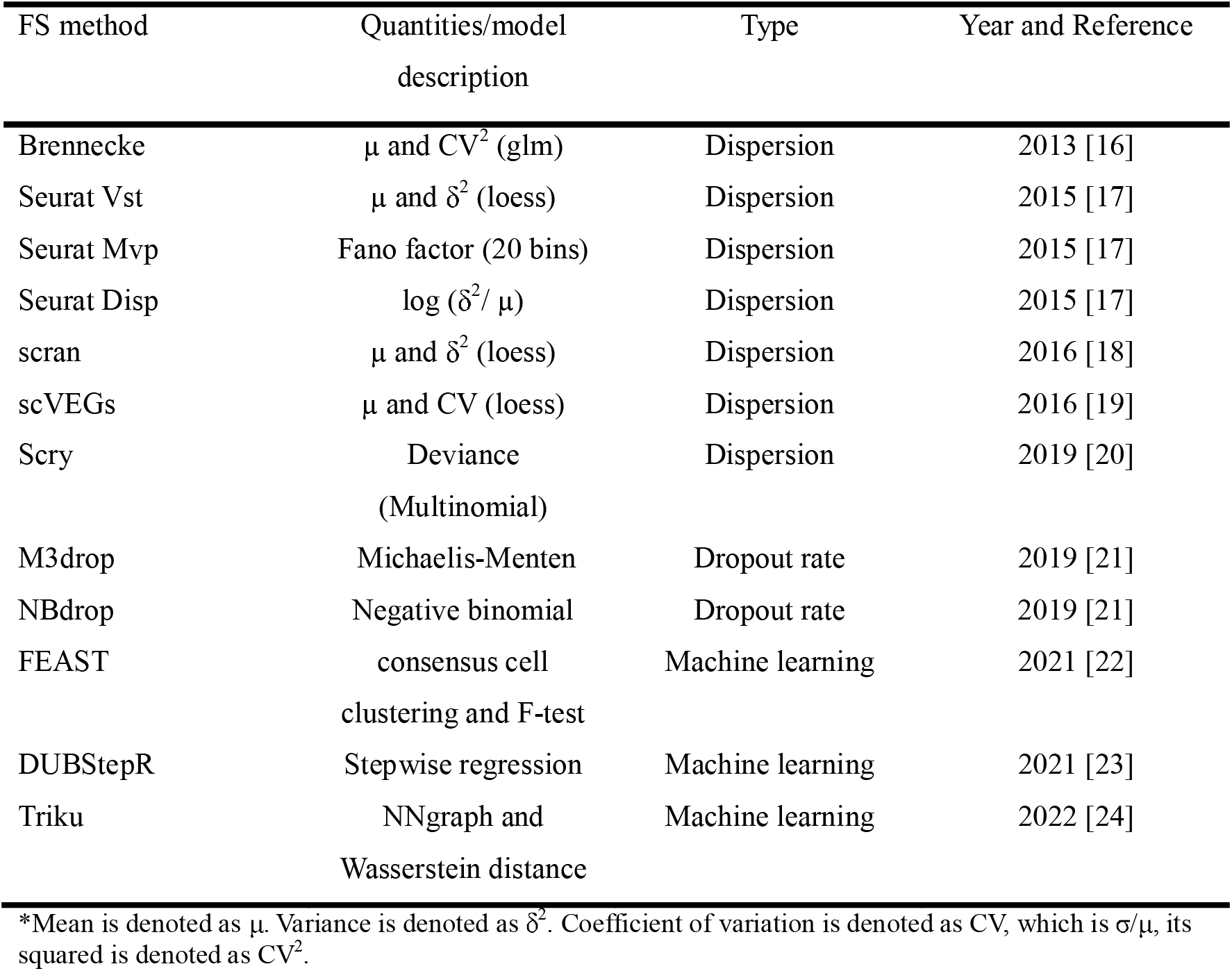
Selected existing feature selection methods in scRNA-seq analysis.

In the case of scran, high variance genes (HVGs) are determined by employing local polynomial regression (loess) on the mean-variance relationship of log-transformed expression values. scVEGs assumes a negative binomial distribution for the mean versus CV relationship and fits this relationship through modified local regression and nonlinear least squares curve fitting. Consequently, parameters of the gene variation model are estimated, enabling the identification of statistically significant variably expressed genes. Within the Seurat tool, three feature selection methods are available. The “vst” method fits a line to the relationship between log variance and log mean using LOESS, similar to scran. The “mvp” method utilizes the Fano factor (variance divided by mean) and categorizes Fano factors into 20 expression mean-based bins. It then normalizes the Fano factors in each bin into z-scores and selects genes accordingly. The “disp” method selects genes with the highest dispersion values. Additionally, Scry calculates a deviance statistic for counts based on a multinomial model assuming a constant rate for each feature. All five methods mentioned above are variance-to-mean approaches that aim to measure dispersion or noise-to-signal ratio. However, they may inadvertently select many low-expression genes due to a high rate of dropouts.

To address this issue, Andrews et al. [21] introduced two new feature selection methods: M3drop and NBdrop, which are based on dropout rates. M3drop is designed for read counts from full-transcript sequencing protocols (e.g., SmartSeq2), while NBdrop is suitable for counts of Unique Molecular Identifiers (UMIs) from tag-based protocols (e.g., 10X Chromium). Both methods operate under the assumption that genes with a higher proportion of zeros than expected could be biologically significant since they are expressed in fewer cells than anticipated. These cells may correspond to specific cell types or states. M3drop fits a Michaelis-Menten model to the relationship between mean expression and dropout rate, while NBdrop employs a negative binomial model for the same purpose.

Recently, novel feature selection techniques for analyzing single-cell RNA sequencing (scRNA-seq) data have emerged, incorporating machine learning and graph-based algorithms. These methods, namely FEAST [22], Triku [23], and DUBStepR [24], differ from the aforementioned model-based approaches as they leverage machine learning techniques. FEAST begins by obtaining a consensus cell clustering and then evaluates the significance of each feature using an F-test, ranking the features based on the F-statistics. Triku identifies genes that are locally overexpressed in neighboring cell groups by examining the count distribution in the vicinity of each cell and comparing it to the expected distribution. The calculation involves determining the Wasserstein distance between the observed and expected distributions, and genes are ranked based on this distance. A larger distance indicates localized expression in a subset of cells with similar transcriptomic profiles. DUBStepR is a stepwise procedure for identifying a core set of genes that strongly reflect coherent expression variation in a dataset. The authors propose a novel graph-based measure for aggregating cells in the feature space, optimizing the number of features based on this measure.

Inspired by these three papers, we recognized the potential value of graph-based machine learning techniques in the field of scRNA-seq data analysis, particularly in the context of feature selection. Consequently, we selected seven existing feature selection methods for comparison in this study: Brennecke, Scry, M3drop, NBdrop, Seurat Vst, Seurat Mvp, and Seurat Disp.

## 3. Datasets

To perform the simulation study, we generated 48 artificial datasets using the SPARsim package [25] in R. SPARsim utilizes a Gamma-Multivariate hypergeometric probabilistic model to create count matrices that closely resemble the distribution of zeros observed in real count data. In a recent evaluation of 16 scRNA-seq simulation methods by Crowell et al. [26], SPARsim produced results that were most similar to real data, followed by Splatter, which is a widely used simulator for scRNA-seq data. Furthermore, SPARsim was highly recommended in a benchmark study by Cao et al. [27] for systematic evaluation of simulation methods for scRNA-seq data. These 48 datasets are divided into two groups: 24 fine-resolution and 24 coarse-resolution datasets. Each dataset consists of 6,000 cells and 15,000 genes, with a total of 10 cell populations. The abundances of these cell populations are as follows: 25%, 20%, 16%, 10%, 8%, 7%, 6%, 4%, 3%, and 1% of the total cells.

To set up the initial parameters, we estimated them using the 10X Genomics example datasets from Zheng et al. [28], specifically the human Jurkat and 293T cells, which are also included as built-in datasets in the SPARsim package. Based on the estimated parameters from 1,718 293T cells, we created 10 different cell groups with the respective abundances mentioned above. In each cell group, we designated 20 genes out of the 15,000 genes as driver genes, which were only upregulated in one type of cell. Therefore, there were a total of 200 driver genes (20 genes per cell group). Additionally, we selected another 1,800 genes to be differential expression (DE) genes, which were either upregulated or downregulated in two or more cell types. This resulted in a total of 2,000 informative genes.

The SPARsim package uses fold change multiplier to control the magnitude of differential expression. A fold change multiplier of 1 indicates no differential expression, while values greater than 1 represent upregulation and values less than 1 represent downregulation. For the upregulated and downregulated DE genes, we generated the fold change multipliers from uniform distributions: Unif (*a_Up_*, *b_Up_*) and Unif (*a_down_*, *b_down_*) respectively.

For the fine-resolution datasets, we defined 24 levels of intensities by selecting the parameters, *a_Up_* ∈ (1.5, 2), *b_Up_* ∈ (2, 2.5), *a_down_* ∈ (0.4, 0.6), *b_down_* ∈ (0.6, 0.8). These four parameters were chosen 24 times in equally spaced intervals within the specified ranges. Similarly, for the coarse-resolution datasets, we set up 24 levels of intensities by selecting parameters *a_Up_* ∈ (2, 3), *b_Up_* ∈ (2.5, 3.5), *a_down_* ∈ (0.1, 0.3), and *b_down_* ∈ (0.3, 0.5). Each dataset in the fine- or coarse-resolution group corresponds to a specific DE fold change multiplier.

Regarding the driver genes, the parameters in the uniform distributions Unif (*a_Up_*, *b_Up_*) were multiplied by 1.2. As the value of *a_Up_* increases and *a_down_* decreases, the differential expression of the genes becomes more pronounced between cell groups, making these genes easier to identify using feature selection methods. Therefore, we considered the 24 datasets with more extreme differences between cell groups as fine-resolution simulated datasets, while the other 24 datasets were coarse-resolution simulated datasets.

In our study, we utilized scRNA-seq datasets provided by 10X Genomics, specifically the datasets of 2,700 peripheral blood mononuclear cells (PBMCs) from the recently published CITE-seq reference dataset by Hao et al. [29]. This reference dataset consisted of 162,000 PBMCs measured using 228 antibodies. The benchmarking dataset included 24 human PBMC datasets, which represented 8 different donors and 3 different batch times. These datasets were manually annotated at three different resolutions: fine, moderate, and coarse annotation. The fine annotation contained 56 cell types, but due to some cell types having zero cells, we decided to focus on the moderate and coarse annotation for our working datasets. The 24 datasets with coarse annotation labels were considered our real-world coarse-resolution datasets. These datasets consisted of 8 different cell types: B cells, CD4 T cells, CD8 T cells, dendritic cells (DCs), monocytes (Mono), natural killer cells (NK cells), Other cells, and Other T cells. For our fine-resolution datasets, we used the moderate resolution labels but restricted them to T cells. This resulted in 24 fine-resolution datasets containing 11 T cell types: CD4 cytotoxic T lymphocytes (CTLs), CD4 naïve T cells, CD4 central memory T cells (TCMs), CD4 effector memory T cells (TEMs), CD8 naïve T cells, CD8 TCMs, CD8 TEMs, double-negative T cells (dnT), gamma-delta T cells (gdT), mucosal-associated invariant T cells (MAIT), and regulatory T cells (Tregs).

To summarize, we had a total of 96 datasets: 24 datasets each for simulation coarse-resolution datasets, simulation fine-resolution datasets, PBMC coarse-resolution datasets, and PBMC fine-resolution datasets.

To determine the ground truth informative gene list for the simulated datasets, we explicitly specified which genes were considered informative, totaling 2,000 genes. Obtaining the ground truth informative gene list for the PBMC datasets, however, was more challenging since we did not have that information available. To address this, we utilized the FindAllMarks() function from the Seurat package to identify highly variable genes (HVGs) for each annotated cell type in the PBMC datasets. This approach allowed us to obtain a semi-ground truth for the PBMC datasets. We set the parameters “return.thresh = 1” and “logfc.threshold = 0” in the FindAllMarks() function to collect all HVGs. Within each cell type, we ordered the genes based on their adjusted p-values in ascending order. If there were fewer than 400 genes with adjusted p-values less than 0.05, we included all of them. However, if there were more than 400 such genes, we selected the top 400 genes based on their adjusted p-values. These collected genes were combined to create a true informative gene list. It should be noted that there might be duplicate genes in this list, specifically when a gene had an adjusted p-value less than 0.05 and was selected in more than two cell types. On average, each PBMC dataset yielded approximately 1,500-2,000 informative genes using this approach. The designation of highly variable genes (HVGs) for the simulated datasets is provided in **Table S1**. Additionally, the summary information for the PBMC datasets, including both coarse- and fine-resolution datasets, can be found in **Table S2**.

## 4. Methods

### 4.1. Mcadet Workflow

Our method can be summarized into five main steps, as depicted in **Figure 1**:

1. Matrix pre-processing: We employ a fuzzy coding approach to transform the data, ensuring it meets the necessary requirements for using MCA, the detailed explanation for this is provided in **Supplementary Methods**. The transformed matrix, denoted as ***P***_n×2p_, where *n* represents the number of cells and *p* represents the number of genes.
2. MCA decomposition: Using the fuzzy coding transformed matrix *P*_nX2p_ obtained in step 1, we calculate the standardized Pearson residual matrix and perform MCA decomposition. The transformed matrix is decomposed into two parts: extrinsic variation *rc^T^* and intrinsic variation 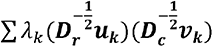. The intrinsic variation is of particular interest as it holds greater biological meaning. It can be further divided into two parts: standard coordinates of rows (cells) 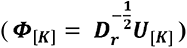 and principal coordinates of columns (genes) (***G_[K]_*** = 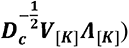, where *K* is the number of principal components we want to keep.
3. Community detection (cells clustering): Cells and genes are plotted in the same K-dimensional space based on *Φ*_[_*_K_*_]_ and *G*_[*K*]_. The Leiden algorithm is applied using a similarity matrix derived from a k-nearest neighbor graph of cells, with the shared nearest neighbor matrix serving as weights.
4. Calculation and ranking of Euclidean distances: After establishing *d* cell communities (clusters), the coordinates of each cluster centroid are computed by averaging the coordinates of cells within the cluster. Using these low-dimensional MCA space coordinates, we calculate the Euclidean distances between each gene and each cell centroid. Based on these distances, we rank the genes within each community, generating a rank list Ω_j_ for each gene: Ω_j_= rd_(j,C1)_,.. .,rd_(j,Cψ)_,.. ., rd_(j,Cd)_1, where rd_(j,Cψ)_ represents the rank of gene J in the Euclidean distances between all genes and the centroid of cell community ψ. The maximum and minimum values from Ω_j_, i.e., *max*_(_Ω_j_), and *min*(Ω_j_), indicate gene variability.
5. Statistical testing: We propose a statistical test to determine whether a gene should be classified as a variable gene. To do this, we define the test statistic as 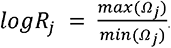. We employ a Monte Carlo simulation approach to calculate p-values. By randomly drawing ranks from 1 to p and obtaining 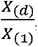, we repeat this process T times (typically a large number) to generate a simulated distribution. Consequently, p-values can be derived from this simulated distribution. All the mathematical derivations and parameter selections are presented in the **Supplementary Methods**.

### 4.2. Evaluation metrics of feature selection performance

In the context of feature selection, the goal is to identify the most relevant features that effectively describe and comprehend the datasets. Specifically, this involves identifying subsets of genes that, when used as inputs in a clustering algorithm, can yield a clustering solution where each cluster represents a potential cell type. To address this, we approached the problem from two perspectives: 1) the accuracy of the selected genes, and 2) the performance of the clustering algorithm.

For assessing the accuracy of the selected genes, we used the Jaccard similarity index as the evaluation metric. The Jaccard similarity index is a statistical measure used to assess the similarity and dissimilarity between two sample sets . It calculates the similarity by dividing the size of the intersection of the sets by the size of their union:

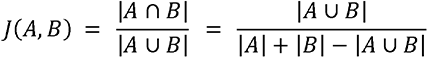

0 ≤ J(A,B) ≤ 1, where J(A,B) close to 1 indicates a higher similarity between the sets *A* and *B*. In our context, the gene sets selected by different feature selection methods represent set *A*, while the (semi-) ground truth of highly variable genes (HVGs) for each dataset, as specified in Section 3, represents set *A*. By comparing the Jaccard similarity index, we can evaluate the accuracy of the selected HVGs.

To assess the clustering performance of different feature selection methods, we initially created reduced-dimensional datasets using the selected informative gene sets. Next, we utilized Principal Component Analysis (PCA) to transform the scRNA-seq data into a lower-dimensional space, specifically reducing the dimensionality to first 15 principal components (PCs). Subsequently, we applied the *k*-means algorithm to cluster the data within this reduced space. We opted for *k*-means due to its effectiveness in low-dimensional spaces and its suitability when the number of cell clusters in both artificial and real-world datasets is known, allowing for straightforward specification of the value of *k*. After applying the k-means algorithm, we obtained the set of clustering denoted as *T*_1_, while the true class is denoted as *T*_2_.

To evaluate the quality of the clustering results, we employed various metrics, including the silhouette score, purity, adjusted Rand index (ARI), normalized mutual information (NMI), and the nearest neighbor graph retention rate (NNgraph retention rate). The first five metrics mentioned are well-established and widely used evaluation metrics for assessing clustering performance. However, the last metric mentioned is a specific evaluation metric devised for this study. These metrics provide insights into different aspects of clustering performance and help us compare the effectiveness of each feature selection method.

The silhouette value is a measure of how well an object is assigned to its own cluster compared to other clusters [30]. It ranges from −1 to 1, with a higher value indicating a better match to its own cluster and a poorer match to neighboring clusters. A higher average silhouette value, known as the silhouette score, indicates that the clustering is more suitable, whereas a lower or negative silhouette score may suggest an insufficient number of clusters or an incorrect clustering configuration. The purity of a cluster is a measure of the proportion of data points belonging to the most frequent class within that cluster [31]. It is calculated by summing the number of data points in each cluster that belong to the most common class and dividing it by the total number of data points *N*. The purity can be defined as follows:

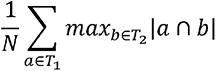

Rand Index is a measure of similarity between two data clusters in statistics, and in particular in data clustering [32]. Normalized Mutual Information is a variation of a measure in information theory known as Mutual Information [33]. Mutual information refers to the amount of information about a distribution that can be derived from a second distribution. It is a good measure for determining the quality of clustering. A major reason that it is usually considered is that it has a comprehensive meaning and can be used to compare two partitions, even when there is a difference in number of clusters. Normalized Mutual Information is given by:

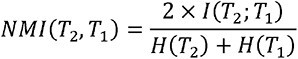

where *H*(·) is the entropy function, *I*(*T*_2_,*T*_1_) = *H*(*T*_2_)–*H*(*T*_2|_T_1_) is the mutual formation between *T*_2_ and *T*_1_.

The NNgraph retention rate is an additional evaluation metric used in this study to assess the performance of feature selection methods. It measures the proportion of nearest neighbors of a cell that belong to the same class. This metric ranges from 0 to 1, where a higher value indicates better clustering performance. Upon a *k* nearest neighbor graph is built by cells in a low-dimensional space using Euclidean distance on first 15 PCs mentioned above, each cell has *k* neighbors, among its *k* neighbors, we define the proportion of that belongs to the same class as it as NNgraph retention rate for single cell:

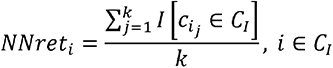

where *c_ij_* is the *j^th^* nearest neighbor cell of cell *i*,*I*[·] is the indicator function, *C_I_* is the class of cell *i*. The NNgraph retention rate for the whole sample of cells is given by:

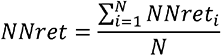

where *N* is the total number of cells. We conducted the analysis using a value of *k*= 30 as the number of neighbors for calculating the NNgraph retention rate.

As mentioned in Section 2, we compared Mcadet with 7 existing feature selection (FS) methods. In addition, we included a “Random” method as a reference for comparison. The details of each method are as follows:

1. Random: Randomly select 2,000 genes from the datasets.
2. Seurat Disp: Uses the FindVariableFeatures() function with “method=disp” from the Seurat package.
3. Seurat Vst: Uses the FindVariableFeatures() function with “method=vst” from the Seurat package, which is equivalent to the scran method described in Section 2.
4. Seurat Mvp: Uses the FindVariableFeatures() function with “method=mvp” from the Seurat package.
5. Brennecke: Uses the BrenneckeGetVariableGenes() function from the M3Drop package.
6. NBdrop: Uses the NBumiFeatureSelectionCombinedDrop() function from the M3Drop R package.
7. M3drop: Uses the M3DropFeatureSelection() function from the M3Drop R package.

## 5. Results

In feature selection, the main goal is to identify the most relevant features that effectively describe and understand datasets. Specifically, this involves finding subsets of genes that, when utilized as inputs in a clustering algorithm, can yield a clustering solution where each cluster represents a potential type of cell. In this study, we conducted a comprehensive evaluation of various existing scRNA-seq feature selection methods. Our evaluation was based on both subjective judgment (qualitative results) and objective evaluation metrics (quantitative results), using a range of synthetic and biological datasets. Detailed information about the datasets and key evaluation metrics can be found in the Section 3 and Section 4. Moreover, we assessed the coherence of the HVG sets selected by different FS methods within each dataset. Furthermore, we performed experiments to compare the proficiency of our method in detecting HVGs in rare cell populations, thus providing additional evidence of its effectiveness. Lastly, we conducted a comparison of the mean expression levels of selected HVGs using different feature selection methods. The purpose of this comparison was to showcase that our proposed method tends to select genes that are highly informative rather than focusing solely on highly or lowly expressed genes.

### 5.1. Qualitative results: UMAP visualization

**Figure 2** illustrates the UMAP visualization of a fine biological scRNA-seq dataset (PBMC dataset pf donor 7 and batch 3) by different FS methods. In this context, the dataset has 11 types of T cells as described in Section 3. To create the working dataset, we extracted HVGs that were selected using different FS methods. Subsequently, the dataset was log-normalized using a scaling factor of 10,000. To visualize the data, we performed UMAP (Uniform Manifold Approximation and Projection) [34] on the first 15 principal components obtained

**Figure 2.**
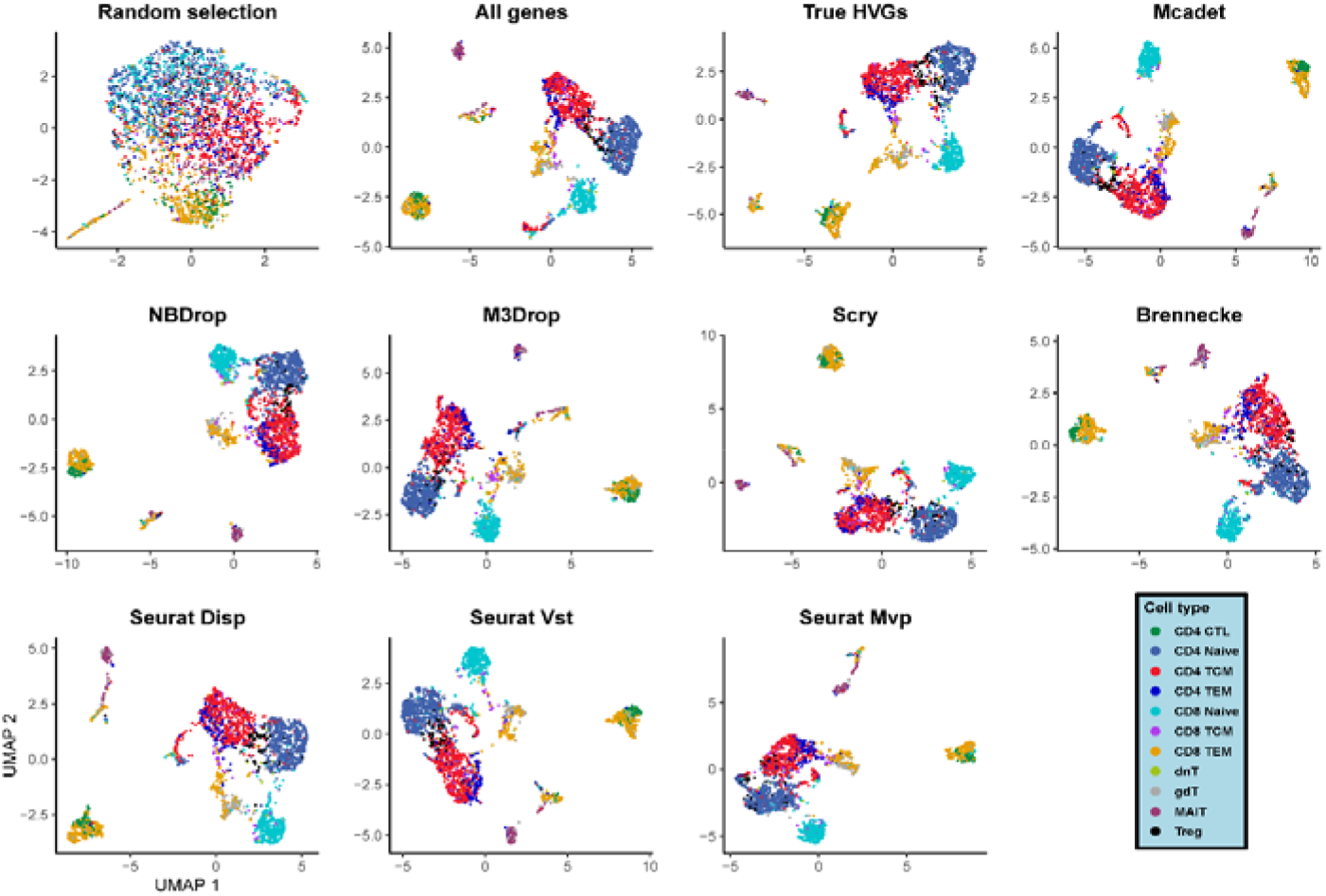
UMAP visualization of a PBMC fine-resolution scRNA-seq dataset. The dataset contains 11 different cell types. The first three plots in the top panel serve as reference plots, emphasizing the significance of feature selection. Subsequently, the remaining plots display the UMAP visualization results obtained from eight different FS methods.

after applying PCA for dimensionality reduction on the log-normalized data. **Figure 2** in the top left plot displays the UMAP visualization of a randomly selected set of 2,000 genes from the entire gene pool. This visualization highlights the importance of feature selection as the separation among cell populations is relatively unsuccessful. The second plot in the top panel demonstrates the UMAP visualization using all the detected genes from the dataset. Although this visualization successfully separates the 11 cell populations, it uses the entire dataset, resulting in longer computation times if no FS performed, especially for downstream analysis of scRNA-seq data. The third plot in the top panel employs a semi-ground truth approach by incorporating the highly variable genes (HVGs) specifically identified from this biological dataset. This approach suggests an ideal separation with acceptable computation time. The remaining plots present the UMAP visualizations generated by each feature selection method using their default parameters. It is important to note that the Seurat Disp and Scry methods can only specify the number of HVGs required, and thus we selected and utilized 2,000 HVGs for both of these methods. Each method exhibits its own unique characteristics. While the visual differences between different feature selection methods may seem minimal at first glance, a closer examination reveals that our proposed method’s UMAP visualization demonstrates the most distinct separation, resembling the true HVGs plot. **Figure S1** illustrates the UMAP visualization of the coarse-resolution PBMC scRNA-seq dataset. Interestingly, even with a random selection of 2,000 genes, a relatively good separation is achieved. This suggests that this particular dataset is relatively easy to differentiate compared to a fine-resolution dataset, where more detailed features are required for accurate separation. In this scenario, the necessity for feature selection is reduced as the dataset inherently exhibits distinguishable patterns without requiring specific feature subsets.

**Figure 3** showcases the UMAP visualization of an artificial fine scRNA-seq dataset (the 12^th^ dataset of 24 simulated datasets) using different FS methods. This dataset was generated using the SPARSim simulator and consists of 10 distinct cell groups. The abundances of these cell groups are as follows: {25%, 20%, 16%, 10%, 8%, 7%, 6%, 4%, 3%, 1%}, as described in Section 3. In this scenario, both random selection and Brennecke feature selection methods fail to successfully separate the 10 cell populations. Furthermore, the visual effects of NBdrop, M3drop, and Seurat Mvp do not resemble the patterns of true highly variable genes (HVGs). The most similar visualization to the true HVGs one is achieved by Mcadet, Scry, and Seurat Disp FS methods. Similarly, in **Figure S2**, the UMAP visualization of the 12^th^ coarse dataset is presented. This observation suggests that when the dataset is inherently easy to differentiate, the choice of FS method may not significantly impact the visualization outcomes. However, the aforementioned results are based on subjective assessments, and it is essential to consider more convincing evaluation metrics.

**Figure 3.**
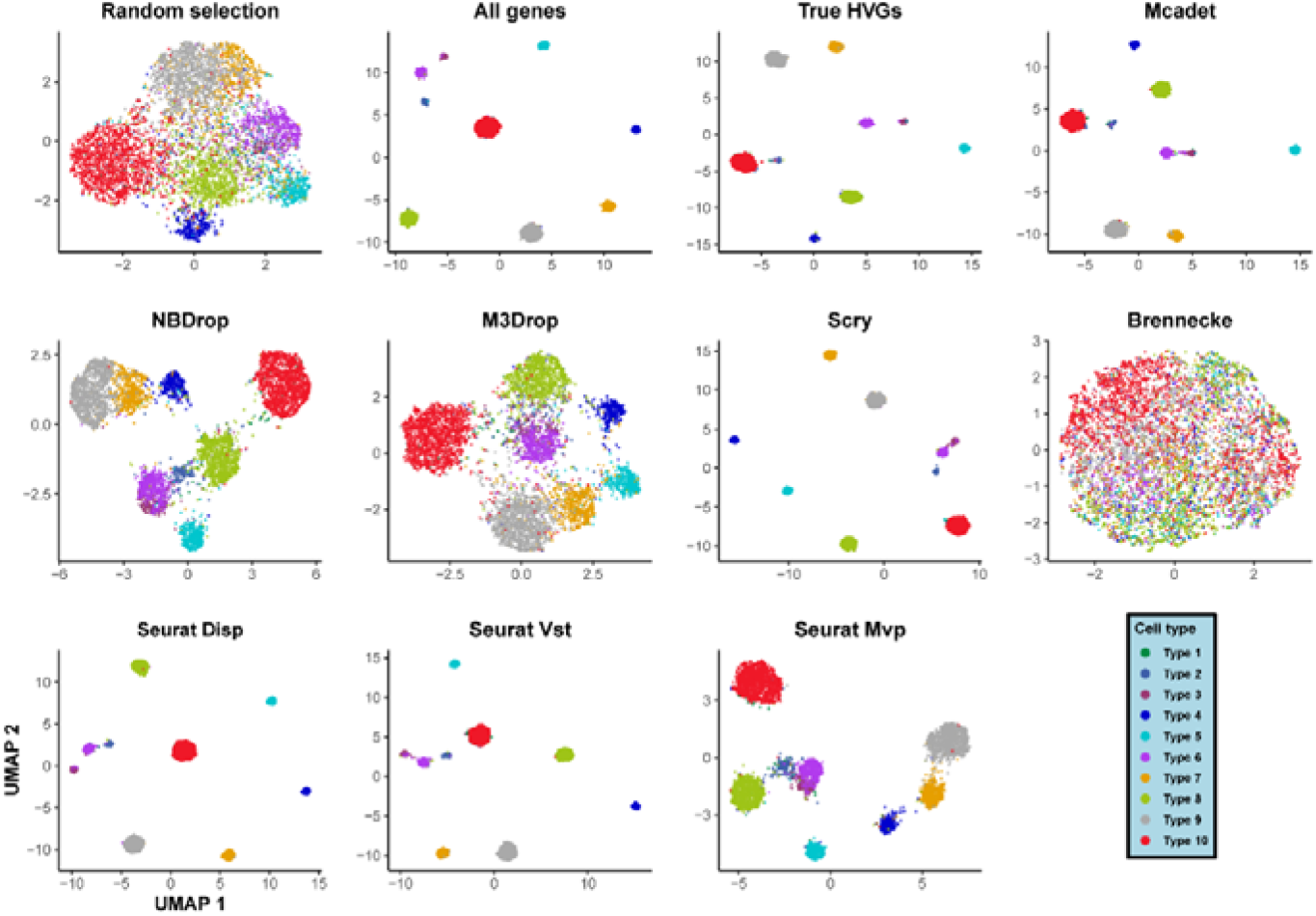
UMAP visualization of a simulated fine-resolution scRNA-seq dataset. The dataset contains 10 different artificial cell types. The first three plots in the top panel serve as reference plots, emphasizing the significance of feature selection. Subsequently, the remaining plots display the UMAP visualization results obtained from eight different FS methods.

### 5.2. Quantitative results: Comparison of evaluation metrics

In our study, we conducted a comparison between our method and other FS methods using six quantitative evaluation metrics: Jaccard similarity index, Silhouette score, Purity, ARI, NMI, and NNgraph retention rate. The Jaccard similarity index primarily evaluates the accuracy of feature selection in selecting highly variable genes (HVGs), while the remaining five metrics assess the clustering performance. More detailed information regarding each evaluation metric can be found in Section 4.

**Figure 4** presents the boxplot of the Jaccard similarity index for different FS methods across various datasets. Each dot represents a dataset from four different dataset types, namely PBMC coarse-resolution, PBMC fine-resolution, simulated coarse-resolution, and simulated fine-resolution datasets. The Jaccard similarity index values were normalized within each type of datasets. **Figure 4** provides compelling evidence that our method achieves a significantly higher mean normalized Jaccard similarity compared to the baseline mean across all four types of datasets (p < 0.001 and 0.0001, t-test). Although our method does not outperform other methods in terms of Jaccard similarity for PBMC coarse-resolution datasets (**Figure 4a**), it emerges as the top-performing method for the other types of datasets (**Figure 4b-4d**), particularly fine-resolution datasets. This outcome highlights a noteworthy enhancement in feature selection accuracy for challenging or difficult-to-differentiate datasets.

**Figure 4.**
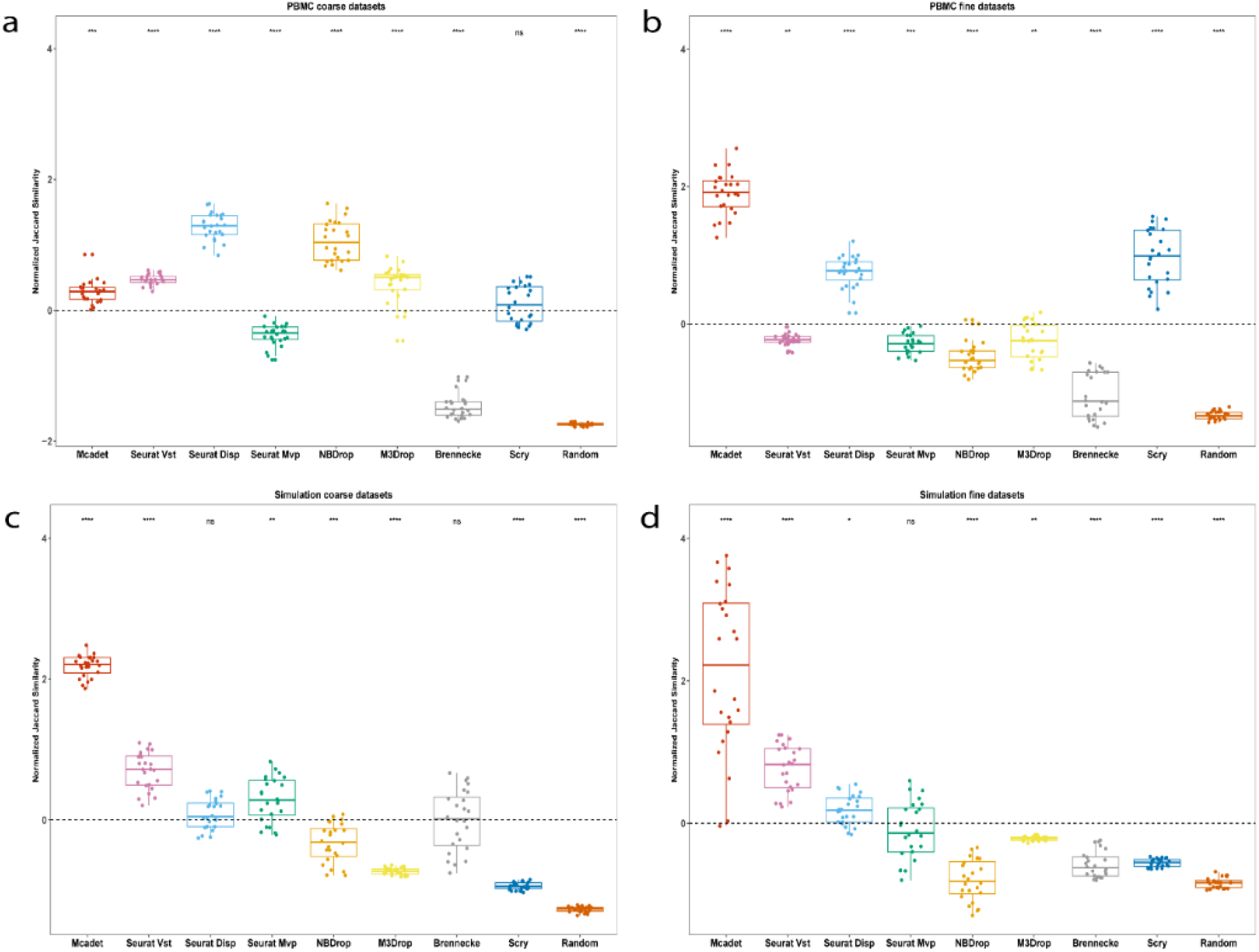
Jaccard Similarity index for comparing feature selection performance on PBMC (a-b) and simulated datasets (c-d). The Jaccard similarity index measures the accuracy of selecting truly highly variable genes (HVGs) by different feature selection (FS) methods. Each dot in the graph represents a dataset, and the dashed horizontal line represents the baseline mean for all methods. The p-values, obtained through t-tests, indicate the significance of the differences between each method and the baseline. “Ns”: non-significance, “*”: p < 0.05, “**”: p < 0.01, “***”: p < 0.001, “****”: p < 0.0001.

**Figure 5-8** exhibit the clustering performance of three evaluation metrics (ARI, purity, and NNgraph retention rate) using boxplots, following the same format as **Figure 4**. The reference method of random selection understandably demonstrates the poorest clustering performance across all three metrics (p<0.0001). For our method, its performance is not significantly different from the baseline mean in the case of coarse-resolution PBMC datasets (p > 0.05). However, when transitioning to fine-resolution datasets, our method consistently outperforms the baseline mean and surpasses other methods (p < 0.0001). These results strongly support our conclusion that our method excels in accurately clustering and distinguishing difficult-to-differentiate datasets. Additionally, we observe that the Seurat Disp and NBdrop methods perform well in coarse-resolution datasets but exhibit lower performance in fine-resolution datasets. Conversely, the Seurat Mvp and Brennecke methods perform poorly in coarse-resolution datasets but show improved performance in fine-resolution datasets, although not as effectively as the Mcadet method. The Silhouette score results are presented in **Figure S3**. In terms of the Silhouette score, our method does not exhibit a significant difference from the baseline mean for both PBMC coarse-resolution and fine-resolution datasets. However, our method demonstrates a significantly higher Silhouette score compared to the baseline mean for the simulation datasets.

**Figure 5.**
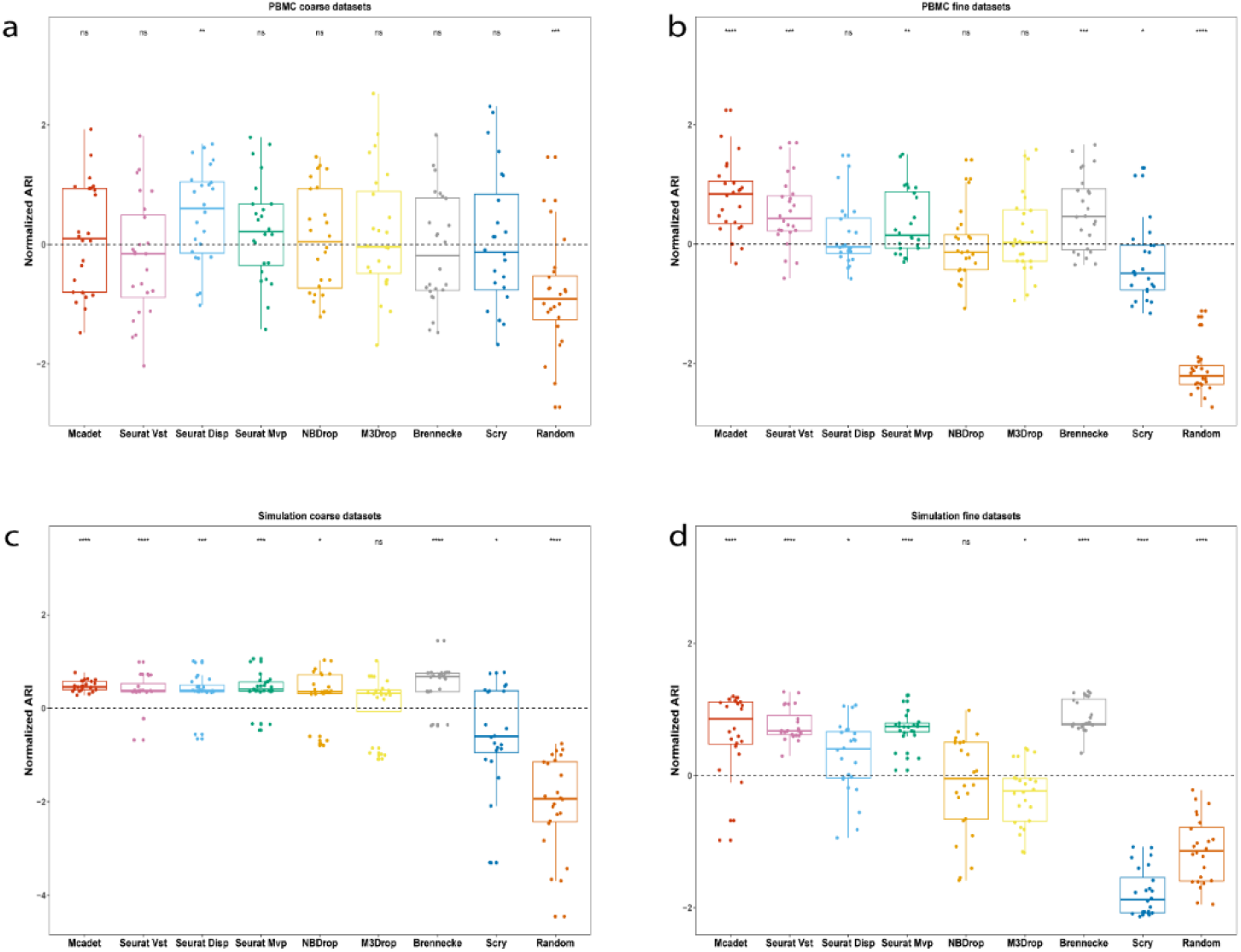
ARI for comparing feature selection performance on PBMC (a-b) and simulated datasets (c-d). ARI is to measure how well the clustering performance with genes selected by different FS methods.

**Figure 6.**
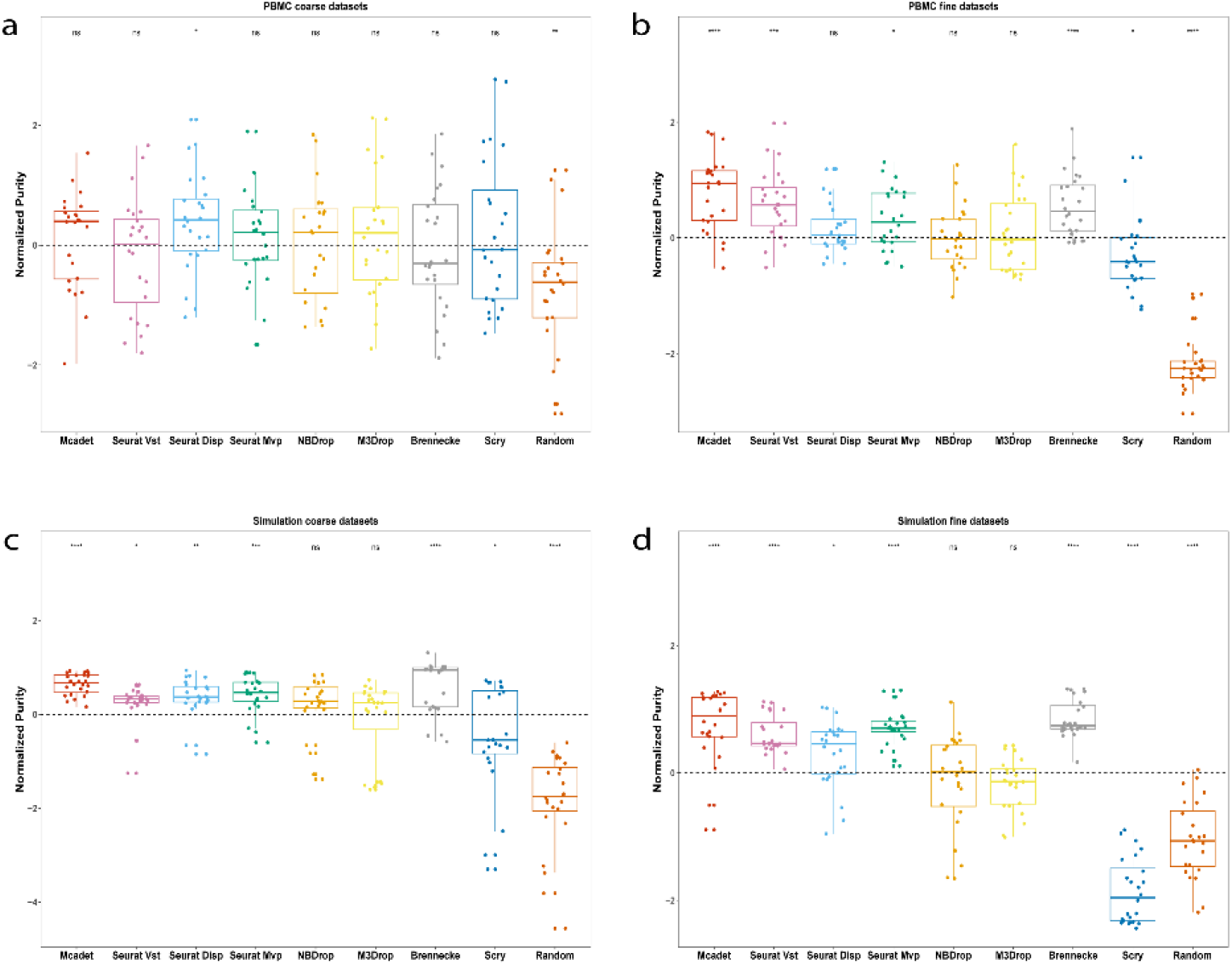
Purity for comparing feature selection performance on PBMC (a-b) and simulated datasets (c-d). Purity is to measure how well the clustering performance with genes selected by different FS methods.

**Figure 7.**
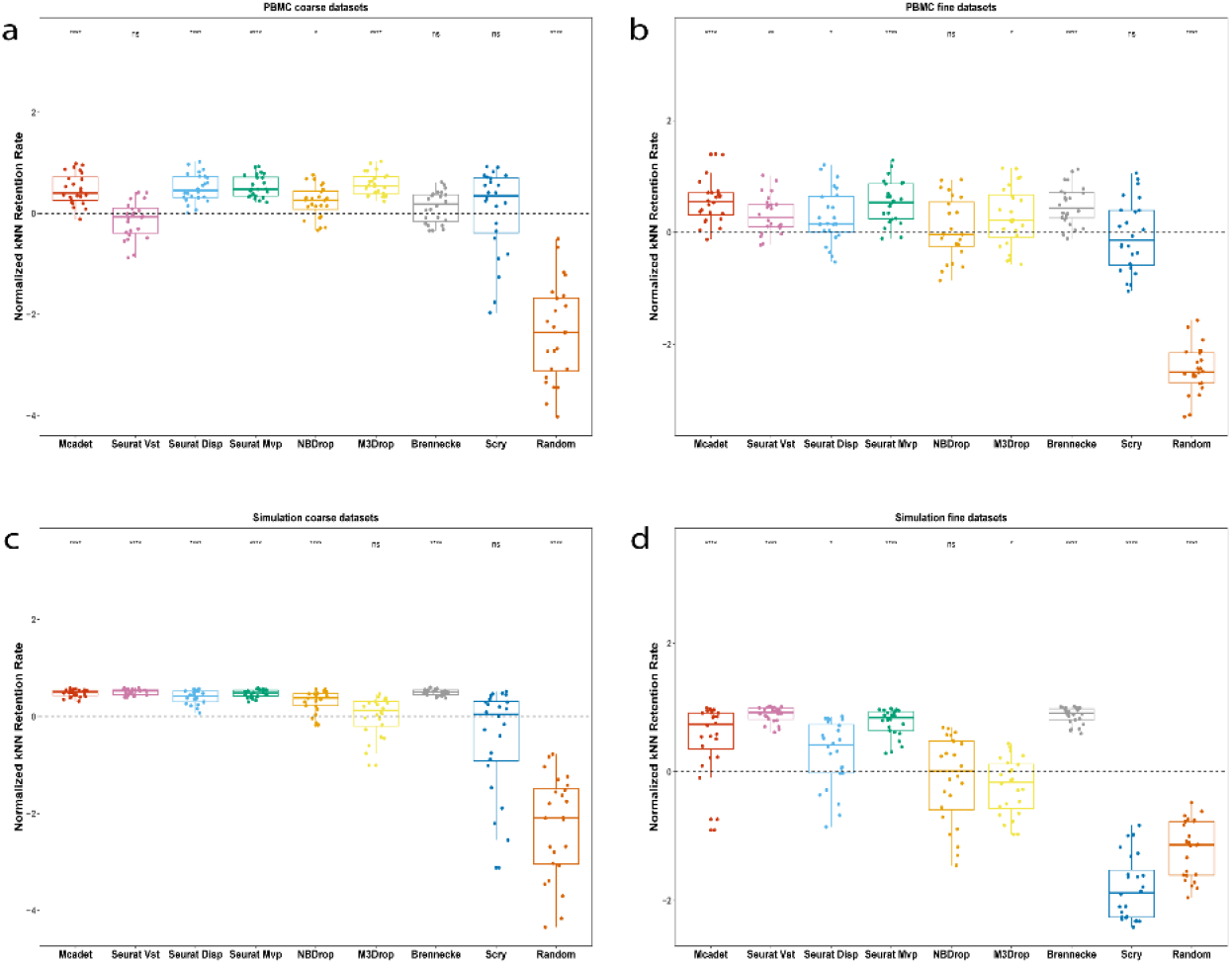
NNgraph retention rate for comparing feature selection performance on PBMC (a-b) and simulated datasets (c-d). NNgraph retention rate is to evaluate the proportion of all the nearest neighbors of a cell belonging to the same cell type.

**Figure 8.**
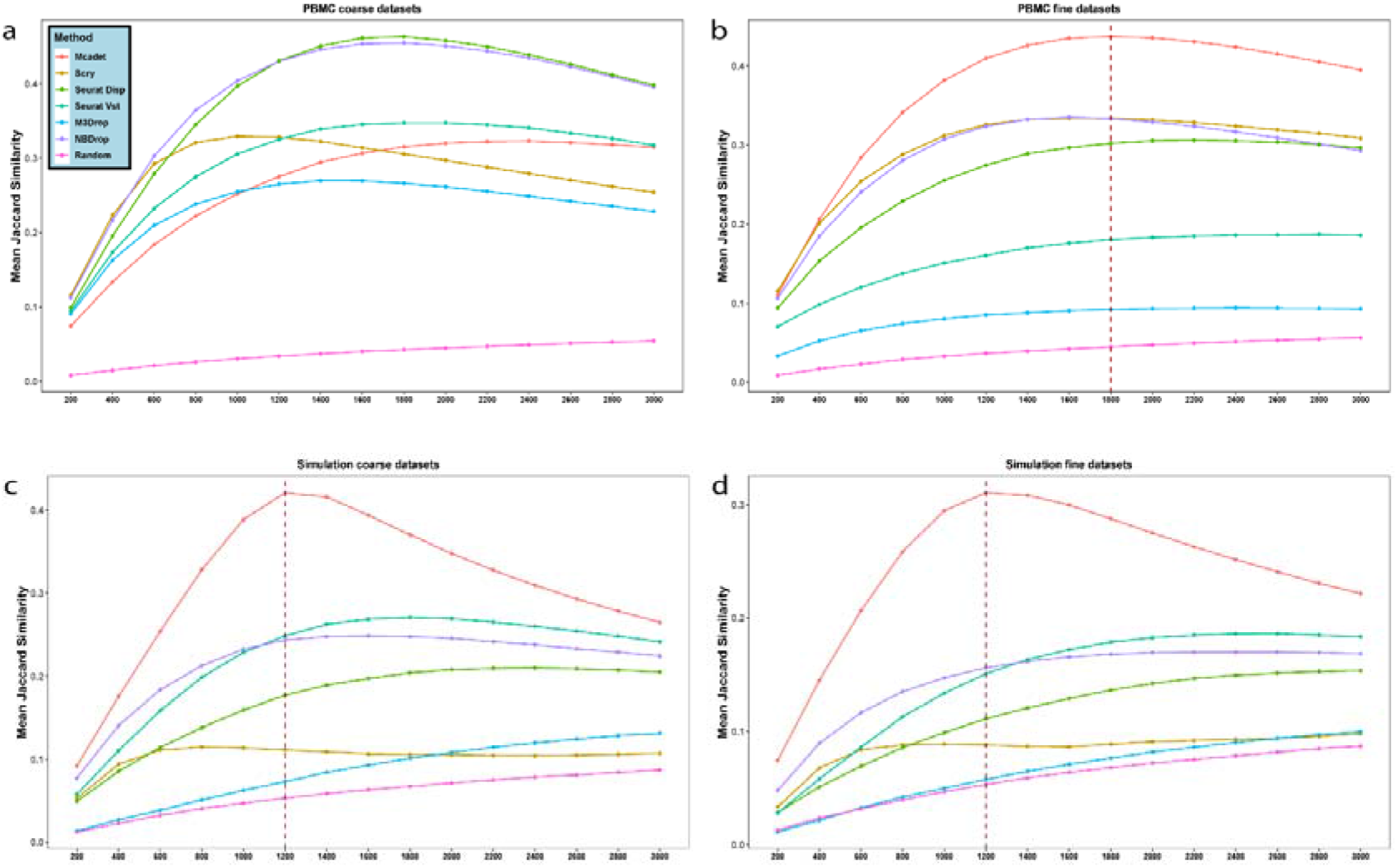
The trend of Jaccard Similarity index as the number of selected genes increases on PBMC (a-b) and simulated datasets (c-d). The number of selected genes range from 300 to 3,000. The Seurat Disp and Scry methods, which only allow for specifying the number of HVGs needed, are excluded from this comparison. The dashed vertical line indicates the number of selected genes that maximizes the Jaccard similarity index when Mcadet exhibits the best performance value.

There is a consensus within the field that the feature selection step in scRNA-seq data analysis should typically involve selecting between 1,000 and 3,000 genes [9]. In a data-driven scenario, the number of informative genes can vary across different datasets, and users may not have prior domain knowledge to determine the exact number of informative genes required. In such cases, our method, along with some existing methods, offers users the opportunity to select informative genes using statistical testing (p-values) or other threshold criteria by default. **Figure 8** presents the trend of mean Jaccard similarity as the number of selected genes increases from 200 to 3,000 across the four types of datasets. The Seurat Disp and Scry methods, which only allow for specifying the number of HVGs needed, are excluded from this comparison. The dashed vertical line indicates the number of selected genes that maximizes the Jaccard similarity index when Mcadet exhibits the best performance value. Observing the graphs, we note that the Jaccard similarity index of our method generally increases until around 1,800 selected genes and then begins to decrease for PBMC fine-resolution datasets. Similarly, for simulation coarse- and fine-resolution datasets, the Jaccard similarity index increases until approximately 1,200 selected genes and then declines. Overall, our method outperforms other methods for these three types of datasets. However, our method does not perform as well as other methods for PBMC fine-resolution datasets, which aligns with our earlier conclusion that our method excels in accurately handling difficult-to-differentiate datasets. Similarly, **Figure S4-S8** depict the trend of clustering performance for different FS methods as the number of selected genes increases. These figures provide additional evidence supporting the effectiveness of our method, particularly in fine-resolution datasets. The clustering performance trends consistently demonstrate that our method performs well when compared to other FS methods in this specific context. We conducted a comparison of the default number of informative genes selected by different feature selection methods, and the results are presented in **Table 2** and **Table 3**. Due to the lack of default parameters for selecting highly variable genes (HVGs) in Seurat Vst, Disp, Scry, and random selection methods, we set the default number of genes to 2,000 for these methods. For the 24 PBMC coarse-resolution datasets, the median number of HVGs selected across all methods is 1,662. Similarly, for the 24 PBMC fine-resolution datasets, the median number of HVGs selected is 1,615. In the case of artificial datasets, the default number of selected genes is fixed at 2,000 for all methods.

**Table 2.** Number of HVGs selected by different FS methods by default. (Simulation datasets)

| Simulated coarse datasets<br>2000 (2000, 2000) |  |  | Simulated fine datasets<br>2000 (2000, 2000) |  |
| --- | --- | --- | --- | --- |
|  | N<br>Median (IQR) | Jaccard Similarity<br>Median (IQR) | N<br>Median (IQR) | Jaccard Similarity<br>Median (IQR) |
| Mcadett | 1356 (1288, 1424) | 0.42 (0.41, 0.43) | 924 (758, 1087) | 0.28 (0.23, 0.34) |
| NBDrop | 506 (473, 547) | 0.17 (0.15, 0.19) | 314 (273, 345) | 0.07 (0.06, 0.09) |
| M3Drop | 6303 (6253, 6365) | 0.13 (0.12, 0.13) | 6420 (6392, 6451) | 0.11 (0.11, 0.12) |
| Brennecke | 1385 (1336, 1434) | 0.20 (0.16, 0.23) | 1231 (1188, 1256) | 0.09 (0.08, 0.10) |
| Seurat Mvp | 682 (640, 713) | 0.23 (0.21, 0.26) | 547 (515, 577) | 0.12 (0.10, 0.14) |
| Seurat Vst | 2000 (2000, 2000) | 0.27 (0.25, 0.29) | 2000 (2000, 2000) | 0.19 (0.16, 0.20) |
| Seurat Disp | 2000 (2000, 2000) | 0.20 (0.19, 0.22) | 2000 (2000, 2000) | 0.14 (0.13, 0.15) |
| Scry | 2000 (2000, 2000) | 0.10 (0.10, 0.11) | 2000 (2000, 2000) | 0.09 (0.09, 0.09) |
| random | 2000 (2000, 2000) | 0.07 (0.07, 0.07) | 2000 (2000, 2000) | 0.07 (0.07, 0.07) |

**Table 3.**
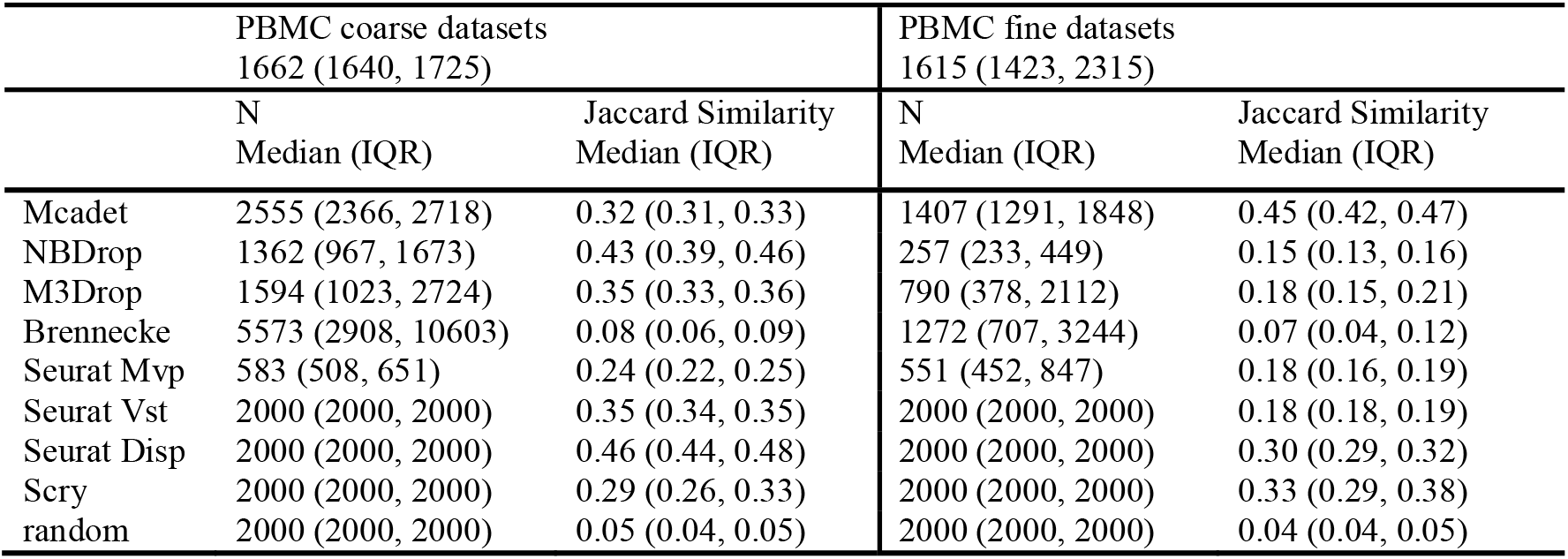
Number of HVGs selected by different FS methods by default. (PBMC datasets)

Our method consistently yields a close match to the true number of HVGs for both PBMC and artificial datasets. Moreover, the corresponding Jaccard index obtained from our method is relatively high compared to the other methods, indicating a better agreement with the ground truth and higher accuracy in selecting informative genes.

### 5.3. Consistency within the same dataset

An additional vital factor to consider when evaluating the effectiveness of a FS method is its consistency within the same sample. Put simply, if a scRNA-seq dataset is randomly divided into two separate datasets by expression value with equal probability, a good FS method will yield similar HVG sets between the two divided datasets, as well as between each divided dataset and the ground truth HVG set. To perform the dataset division, we employed a random splitting method, evenly dividing the datasets into two subsets. The concept of this splitting technique draws inspiration from a method known as count splitting [35]. This was achieved by generating *X*_(1),ij_ ∼ *Binom*(*x*_ij_,*ε*), where *x_ij_*represents the expression value of gene *j*in cell *i* from the original scRNA-seq dataset. Here, ε = 0.5 denotes the probability. Consequently, *x*_(1),ij_ corresponds to the expression value of gene *j*in cell *i* within the first split dataset. The expression value of gene *j*in cell *i* within the second split dataset, denoted as *x*_(2),ij_, is calculated as *x*_ij_ - *x*_(1),ij_ .

**Figure 9** depicts the consistency outcomes for four different types of datasets. Each bar represents the mean Jaccard similarity index of 24 datasets within each data type by each FS method. The error bar signifies the standard error of the Jaccard similarity index. For each FS method, there are three bars. The first (left) bar indicates the mean Jaccard similarity between two separate datasets. The second bar represents the mean Jaccard similarity between the first split dataset and the true HVG set. Similarly, the third bar denotes the mean Jaccard similarity between the second split dataset and the true HVG set. The results have been normalized within the respective data type, with the mean set to 0. Any bar positioned above the blue horizontal line at 0 indicates a performance above the average within the respective group. Conversely, any bar below the blue horizontal line at 0 suggests a performance below the average within the group. In terms of consistency performance between two separate datasets, Mcadet shows an average performance in first three data types (**Figure 9a-9c**). On the other hand, the Scry, NBDrop, and M3Drop methods exhibit the best consistency performance between two separate datasets. As a reference method, random selection demonstrates relatively poorer performance. However, when we consider the second and third bars, Mcadet surpasses other methods. This indicates that the gene sets selected by Mcadet using either the splitting of one or two datasets exhibit a strong correlation with the true HVG set.

**Figure 9.**
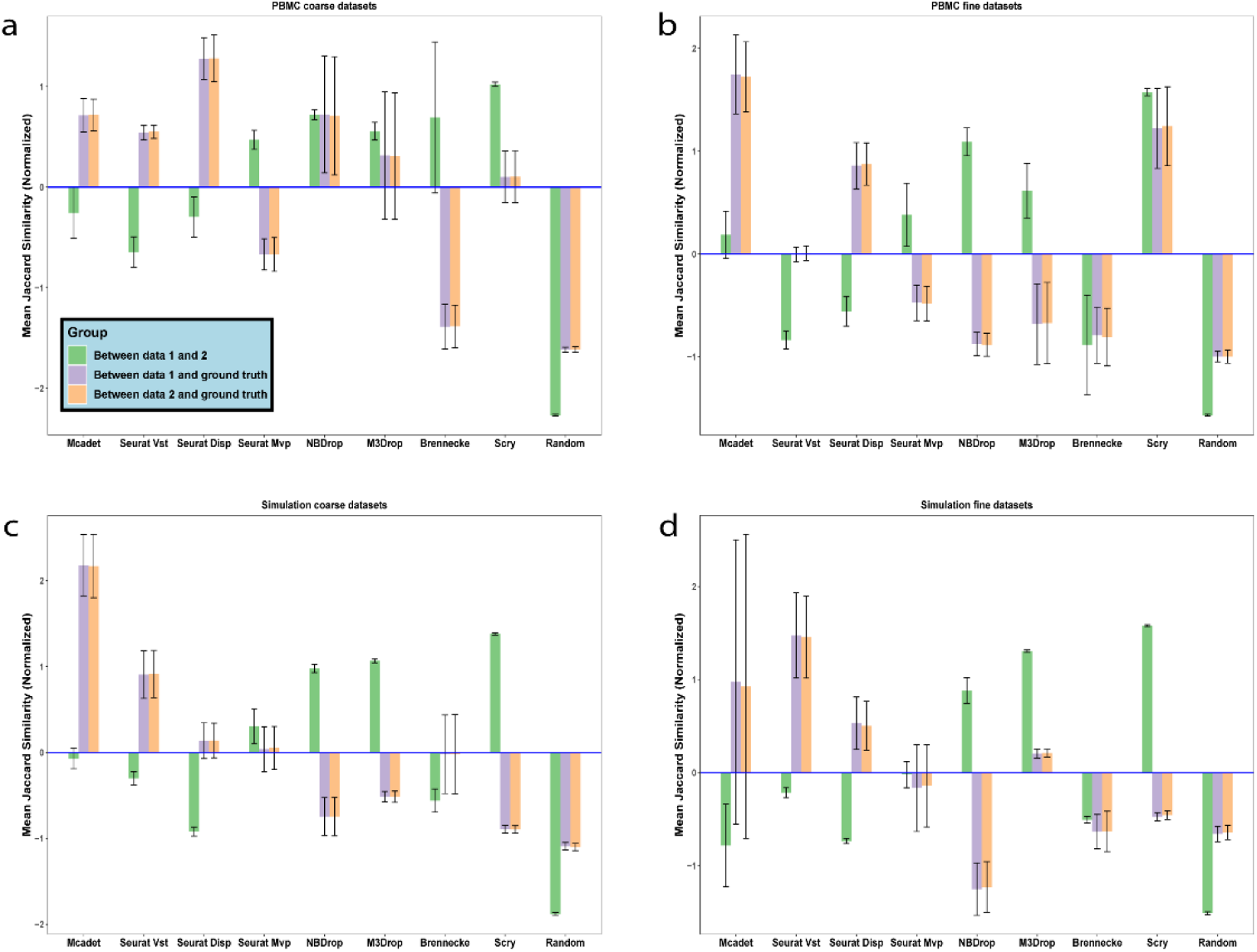
Jaccard similarity index for assessing the consistency on PBMC (a-b) and simulated datasets (c- d). Left: Mean similarity between separate datasets. Middle: Mean similarity between first split dataset and true HVG set. Right: Mean similarity between second split dataset and true HVG set. Results normalized within each data type group, mean set to 0.

This result provides a valuable insight for the application of the Mcadet method: Users can follow our approach by splitting the dataset as described and applying Mcadet to both datasets. Subsequently, they can combine the selected HVGs from both datasets to form the final HVG set. Furthermore, the second and third bars highlight that if the gene expression is artificially reduced (in the case of lowly expressed genes), our method outperforms others in identifying lowly expressed but highly variable genes in such scenarios.

### 5.4. Rare cell analysis

To evaluate the performance of the feature selection (FS) method in selecting highly variable genes (HVGs) for rare cell populations, we incorporated predefined rare cell populations in each dataset. In the case of the 48 simulation datasets (comprising both coarse- and fine-resolution datasets), cell type 1 was specifically designated as the rare cell type (as shown in **Table S1**). For each dataset, cell type 1 consisted of only 60 cells out of a total of 6,000 cells. Additionally, 620 genes were predetermined as HVGs for cell type 1. In the case of the 24 coarse-resolution PBMC datasets, we deliberately selected 60 B cells from each original dataset to establish B cells as the rare cell type. Subsequently, feature selection was performed on these datasets. The true HVGs for B cells were obtained using the “FindAllMarkers(cluster = ’B’)” function in the Seurat package. As for the remaining 24 fine-resolution PBMC datasets, dnT cells were identified as the rare cell type. The true HVGs for dnT cells were obtained using the “FindAllMarkers(cluster = ’dnT’)” function in the Seurat package. The evaluation metric used here is the proportion of true highly HVGs retained. This is calculated by taking the intersection between the gene set selected by the FS method and the true HVG set and dividing it by the total number of genes in the true HVG set. **Figure 10** illustrates the results of the rare cell analysis, similar to **Figure 9**. Each bar represents the mean proportion of HVGs for each FS method, while the blue dashed line represents the baseline mean. Overall, our method consistently ranks among the top three in all four data types, indicating its excellent performance in identifying informative genes for rare cell populations. The Brennecke method also performs well; however, the large standard error suggests its instability in accurately identifying HVGs for rare cells.

**Figure 10.**
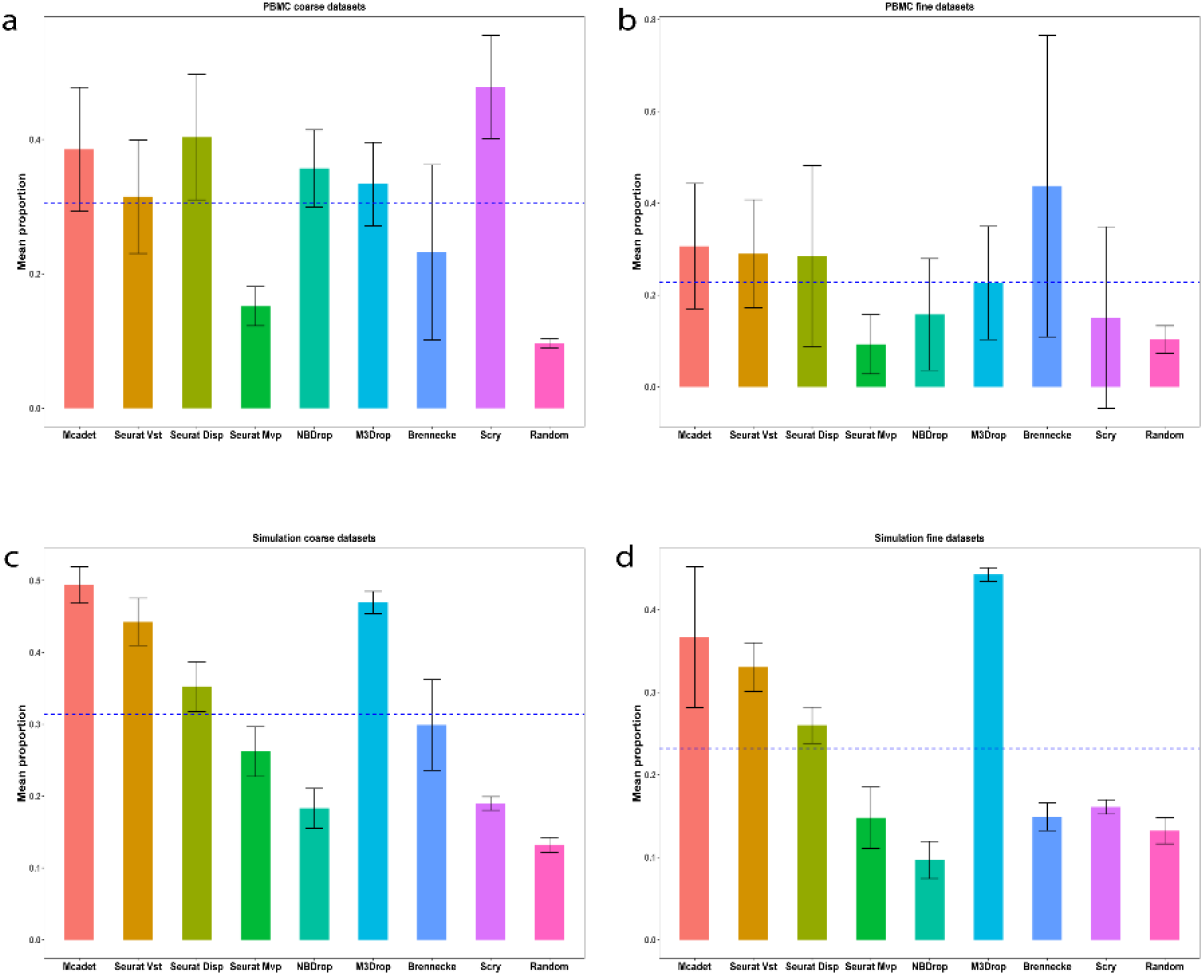
Retention proportion: Performance evaluation on rare cell populations on PBMC (a-b) and simulated datasets (c-d). The blue dashed lines represents the baseline mean within each data type.

### 5.5. Expression analysis

The objective of feature selection is to identify and choose highly variable or highly informative genes. These genes can encompass both highly and lowly expressed genes. It is important to note that certain lowly expressed genes can still hold significant importance in terms of information content. Hence, a good feature selection method should have the ability to capture both highly and lowly expressed genes, rather than being biased towards selecting only highly expressed genes or exclusively focusing on lowly expressed genes. Therefore, it is crucial to consider the density of gene expression for the selected HVGs across different FS methods. **Figure 11** presents the density plot of the log mean expression of the pooled selected genes for each FS method in each data type. The dashed vertical blue line represents the mean for each distribution. It is observed that Mcadet exhibits a unimodal distribution with a normal-like pattern and a moderate mean. Conversely, NBdrop and Scry tend to favor selecting highly expressed genes. Particularly, Scry shows a high concentration of genes with a specific mean expression value. On the other hand, Brennecke tends to select lowly expressed genes. Furthermore, the log mean gene expression density for Seurat Disp and Vst appears to exhibit a bimodal distribution, which is similar to random selection. The result indicates that our method is capable of selecting both highly expressed genes and lowly expressed genes without exhibiting any bias or preference. It demonstrates a balanced approach in capturing genes across a wide range of expression levels.

**Figure 11.**
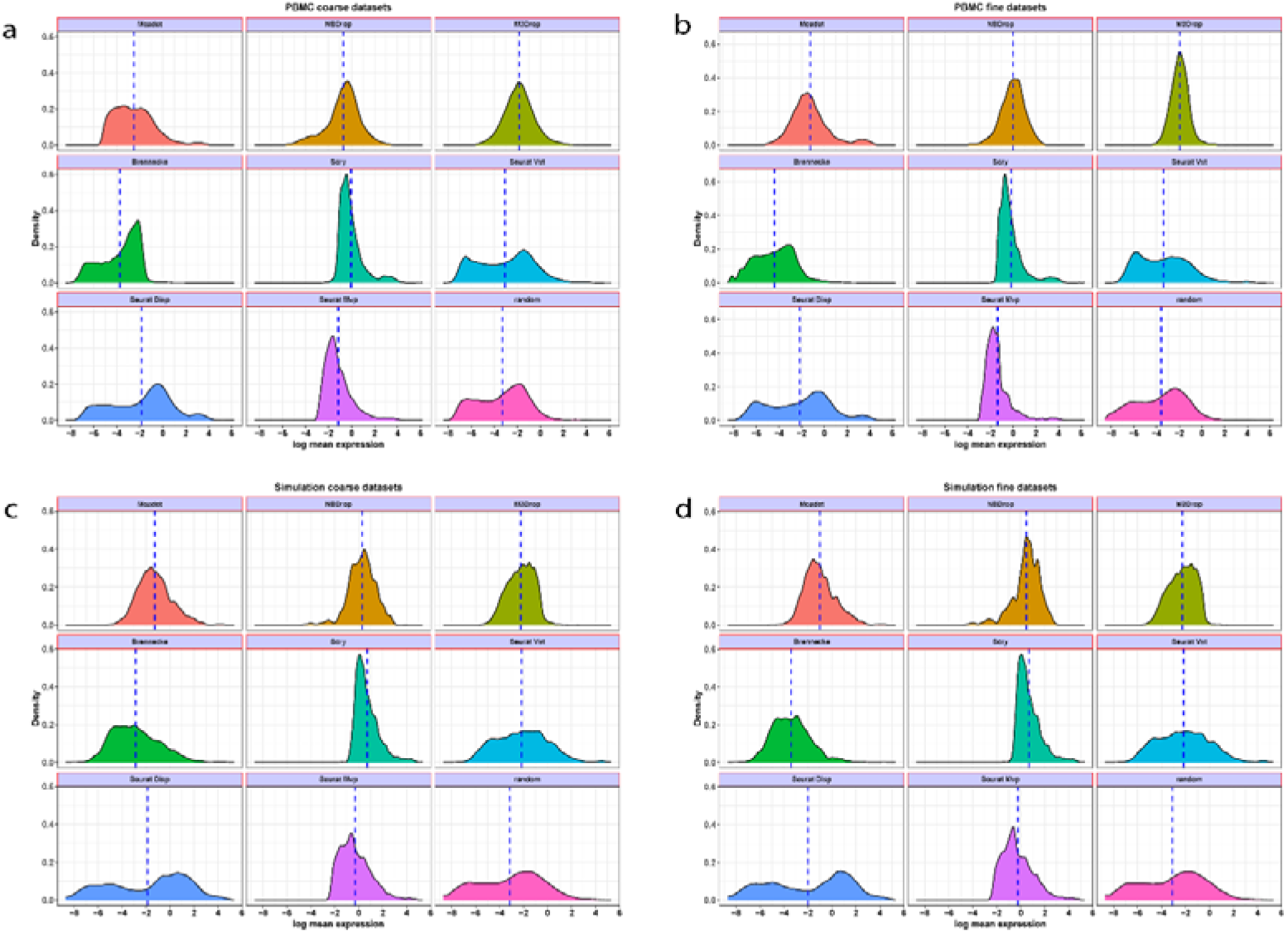
Density of log mean expression of selected genes by FS methods on PBMC (a-b) and simulated datasets (c-d). The Blue dashed lines stand for the mean for each density.

## 6. Discussion

FS methods are integral to scRNA-seq analysis pipelines, as they facilitate the acquisition of a reduced-dimensional dataset that captures crucial information, promoting the interpretation and understanding of the underlying biological processes. In the context of scRNA-seq clustering, the careful selection of a suitable feature set prior to clustering is pivotal, as it directly impacts the quality of the resulting clusters. It is crucial to avoid the inclusion of non- informative genes or the neglect of marker genes, as these factors can significantly compromise clustering accuracy. It is essential to acknowledge that individual FS methods operate based on specific assumptions pertaining to the characteristics that determine the relevance of genes. Certain FS methods adopt gene dispersion as a criterion, postulating that the variability in gene expression is indicative of its biological significance. Conversely, other FS methods, such as NBdrop and M3drop, identify genes with a higher-than-expected proportion of zero-counts, derived from a null distribution, as more informative. Moreover, recent advancements have introduced FS methods that leverage graph-based and clustering machine learning algorithms, such as FEAST and Triku, for feature selection in scRNA-seq data analysis. Despite the extensive development and widespread adoption of various FS methods tailored for scRNA-seq data, there is still a need for exploration, investigation, and proposition of new FS methods adapted to address specific challenges. For instance, certain difficult-to-distinguish datasets or rare cell populations necessitate specialized FS methods that have yet to be thoroughly explored or developed. Further research in this area holds promise for enhancing scRNA-seq analysis and uncovering novel insights into cellular heterogeneity and biology.

In this study, we present Mcadet, a novel FS method designed specifically for UMI scRNA-seq data. Mcadet leverages MCA and community detection techniques to address the challenges posed by difficult-to-differentiate datasets and datasets containing rare cell populations. Using a diverse collection of 96 datasets comprising both simulated and real-world PBMC datasets, we evaluated the performance of Mcadet in identifying highly informative genes. Our assessment considered various evaluation metrics for accuracy and clustering performance. Notably, Mcadet demonstrated excellent performance on datasets with fine resolution, which are often challenging to differentiate. It also exhibited outstanding ability in detecting HVGs associated with rare cell populations. Furthermore, Mcadet successfully determined an appropriate number of HVGs, and in scenarios involving artificially reduced gene expression, our method outperformed other approaches in identifying genes that exhibit both low expression levels and high variability. Particularly Mcadet demonstrated a balanced approach in selecting genes across a wide range of expression levels, without exhibiting any bias or preference. Its ability to capture highly expressed as well as lowly expressed genes reinforces its comprehensive nature.

Our method comprises five major steps: (1). Matrix pre-processing. (2). MCA decomposition. (3). Community detection for cell clustering. (4). Computation and ranking of Euclidean distances. (5). Statistical testing. The efficacy of Mcadet in identifying highly informative genes is primarily attributed to Step (2) and (5), where the MCA decomposition isolates intrinsic variation, which carries greater biological significance, and a novel statistical testing approach is proposed to select a relatively suitable number of HVGs. The notable advantage of Mcadet lies in Step (3), where the application of community detection with varied resolution parameters facilitates robust cell population clustering based on the reduced-dimensional intrinsic biological variation. Consequently, Mcadet exhibits exceptional performance in difficult-to-differentiate datasets and datasets containing rare cell populations. This flexibility enables the accurate detection of fine-resolution patterns and rare cell populations within the data, enhancing the analytical capabilities of Mcadet in challenging scRNA-seq scenarios.

While CA can also isolate intrinsic variation similar to what we achieve in step (2), the use of MCA provides certain advantages. The decision to utilize MCA instead of CA in our approach, stems from the following potential rationale. When applying CA directly, “cells” or “genes” are treated as single categorical variables, with each gene within the “genes” variable regarded as a class or category, and likewise for each cell. CA is designed for visualizing and analyzing the relationship between two categorical variables. However, it is important to acknowledge that genes can exhibit correlation or association, such as gene co-expression, indicating functional relationships [36]. Therefore, it is more appropriate to consider each gene as a variable rather than a category within a single variable, both from a biological and statistical standpoint. Similarly, cells can interact with their neighboring cells [37], suggesting that viewing each cell as a subject rather than a category within the same variable is preferable. In this scenario, where each gene is treated as a distinct categorical variable, the number of variables exceeds two (*p* > 2), making MCA a more suitable choice compared to CA. MCA enables the analysis and visualization of relationships among multiple categorical variables, providing a more comprehensive perspective on the complex structure of scRNA-seq data. While the Cell-ID paper does not explicitly mention the reasons for using MCA instead of CA, this potential rationale aligns with the need to consider genes as separate variables and cells as distinct subjects, leading to the adoption of MCA in our methodology.

To address the absence of categories within each gene variable, we employ fuzzy coding technique in our step (1). Unlike one-hot encoding or crisp coding that assigns only 0 or 1 values, pseudo categories are categorical variables represented in fractional form. This fuzzy coding approach, as demonstrated by Aşan et al. [38] and Greenacre [39], ensures that no information is lost during data transformation and allows for capturing non-linear relationships between variables. By employing fuzzy coding, a variable can be divided into *d*+ 1 intervals by defining *d* cut points. However, it is important to note that this expansion of data into a higher-dimensional space can result in longer computation times for matrix decomposition. For example, if each gene is classified into four categories, a sample with 15,000 genes would expand to 60,000 gene categories. This significant increase in the number of categories presents challenges in terms of computation efficiency and downstream feature selection. Therefore, we adopt the “doubling” technique in correspondence analysis [40, 41]. In this approach, each variable is divided into two categories, namely *g*_+_ and *g*_-_. Here, *g*_+_ represents the presence of gene expression, while *g*_-_ indicates the absence. Only the *g*_+_ categories are extracted for further downstream analysis, reducing the dimensionality, and addressing the issue of selecting gene categories for feature selection.

In step (4) of our approach, the calculation of Euclidean distance involves projecting all genes (*g*_+_), and cells onto the same space, referred to as a biplot. Biplots, originally introduced by Gabriel, K.R., are a type of exploratory graph widely used in statistics. They represent the rows and columns of a data matrix as points or vectors in a low-dimensional Euclidean space. While biplots are typically visualized in two or three dimensions for ease of interpretation, they can be extended to higher-dimensional spaces for computational or analytical purposes. In this low-dimensional space, the proximity, as measured by Euclidean distance, between a column (e.g., gene) and a row (e.g., cell) indicates the degree of specificity of the gene to the cell. For example, if a gene *j*exhibits extreme proximity to cell cluster *A* (i.e., ranking first in distances between all genes and centroid *A*), but is distant from cluster *B*. (i.e., ranking last in distances between all genes and centroid *B*), gene *j* can be considered a candidate HVG that provides informative discriminatory power for distinguishing different cell types or cell states.

In our approach, the task of cell clustering is performed in an unsupervised manner, aiming to discover latent cell types or states within the dataset. While various existing clustering algorithms, such as k-means, DBSCAN, and Gaussian mixture models, can be utilized, they often face limitations when applied to high-dimensional and sparse scRNA-seq data. For instance, k-means requires the number of clusters to be pre-specified, which is challenging for unknown samples. As a result, we turn to graph-based community detection algorithms. One drawback of community detection algorithms is their dependence on the resolution parameter, which influences the clustering outcome. Different resolution values can lead to distinct cluster assignments, making the partitioning result less robust. However, in our context, it is not necessary to precisely determine the exact number of cell types present in the sample. This is due to several reasons: (1). After clustering, the selection of features is based on a novel metric, the log-ratio between the maximum and minimum rank, which is not sensitive to the performance of a specific clustering algorithm. (2). The clustering step in our feature selection workflow aims to enhance the distinguishability of distances between genes and cells, simplifying the identification of informative features. It is not intended to determine the exact number of cell types present, as additional clustering techniques can be applied after the feature selection process. (3). To obtain a more robust clustering result, we consider a range of resolution values and evaluate the overall test statistic. (4). The true partitioning of cell populations can be dynamic, exhibiting variations in fine and coarse resolutions. Therefore, we employ the Leiden algorithm in our feature selection workflow to address this aspect.

However, it is essential to acknowledge the limitations of Mcadet. One limitation is the running time of the method, as demonstrated in **Figure S9**. The experiments were conducted on a desktop with an AMD Ryzen 9 7950X 16-Core Processor and 64 GB RAM. It was observed that the running time of Mcadet increases linearly with the number of genes or cells in the dataset. In comparison to other FS methods, Mcadet exhibits significantly longer running times. This extended duration can be attributed to the inclusion of computationally intensive machine learning steps within our framework, which are absent in other compared FS methods. The second limitation of our approach is the current reliance on the Leiden algorithm for cell clustering. While it is currently the optimal choice for our proposed framework, advancements in clustering or network analysis may lead to the emergence of new and powerful algorithms that could replace Leiden in the future. A third limitation is that although we used 96 datasets, including 48 real-world datasets, all of them are PBMC datasets. In future work, it would be beneficial to include datasets from other cell types or biological systems to further validate the performance and generalizability of Mcadet. Furthermore, it is important to note that we did not consider all available FS methods, such as recently published machine learning-based approaches. Including these methods in comparative evaluations would provide a more comprehensive analysis of Mcadet’s performance and its position among other state-of-the-art techniques. Lastly, we acknowledge that we did not extensively explore optimal parameter settings for Mcadet and instead relied on default parameter configurations. Future investigations should consider conducting parameter optimization to further refine the performance of the method and explore its sensitivity to different parameter choices.

## Code Availability

The Mcadet function and all the R files for implementing the analysis can be accessed on the following GitHub repository: https://github.com/SaishiCui/Mcadet.

## Data Availability

Both the SPARSim simulation datasets and the real-world PBMC working datasets were generated using an R file available at https://github.com/SaishiCui/Mcadet.

Details about PBMC datasets can be found in Section 3. The original real-world PBMC datasets can be obtained from: https://satijalab.org/seurat/articles/multimodal_reference_mapping.html

## Supplementary material

Supplementary material is available for this article online.

## Author Contributions

SC designed the Mcadet algorithm, with assistance from SS and IFZ. SC implemented and performed benchmarking analyses, with assistance from SS and IFZ. SC wrote the manuscript.

## Acknowledgements.

The authors would like to acknowledge Sina Nassiri and Issa Zakeri for their mentorship and guidance, and for critical feedback and discussions.

---

**Table S1.** Highly variable gene designation for simulated datasets.

| Cell population<br>(N, %) | Driver genes<br>designation | HVGs designation<br>(Driver gene excluded) | Number of HVGs (N, %)<br>(Driver gene included) |
| --- | --- | --- | --- |
| Type 1 (60, 1) | Gene 1-20 | Gene 201-400, 801-1000, 1401-1600 | 620 (4.1) |
| Type 2 (180, 3) | Gene 21-40 | Gene 401-600, 801-1000, 1201-1400, 1601-1800 | 820 (5.5) |
| Type 3 (240, 4) | Gene 41-60 | Gene 201-600, 1401-1600, 1801-2000 | 820 (5.5) |
| Type 4 (360, 6) | Gene 61-80 | Gene 401-600, 1001-1600, 1801-2000 | 1000 (6.7) |
| Type 5 (420, 7) | Gene 81-100 | Gene 201-400, 601-800, 1601-2000 | 820 (5.5) |
| Type 6 (480, 8) | Gene 101-120 | Gene 401-600, 1001-1200, 1401-1600, 1801-2000 | 820 (5.5) |
| Type 7 (600, 10) | Gene 121-140 | Gene 201-400, 601-800, 1201-1400 | 620 (4.1) |
| Type 8 (960, 16) | Gene 141-160 | Gene 601-800, 1001-1200, 1601-1800 | 620 (4.1) |
| Type 9 (1200, 20) | Gene 161-180 | Gene 201-400, 1201-1400, 1601-2000 | 820 (5.5) |
| Type 10 (1500, 25) | Gene 181-200 | Gene 801-1200, 1401-1800 | 820 (5.5) |

**Table S2.**
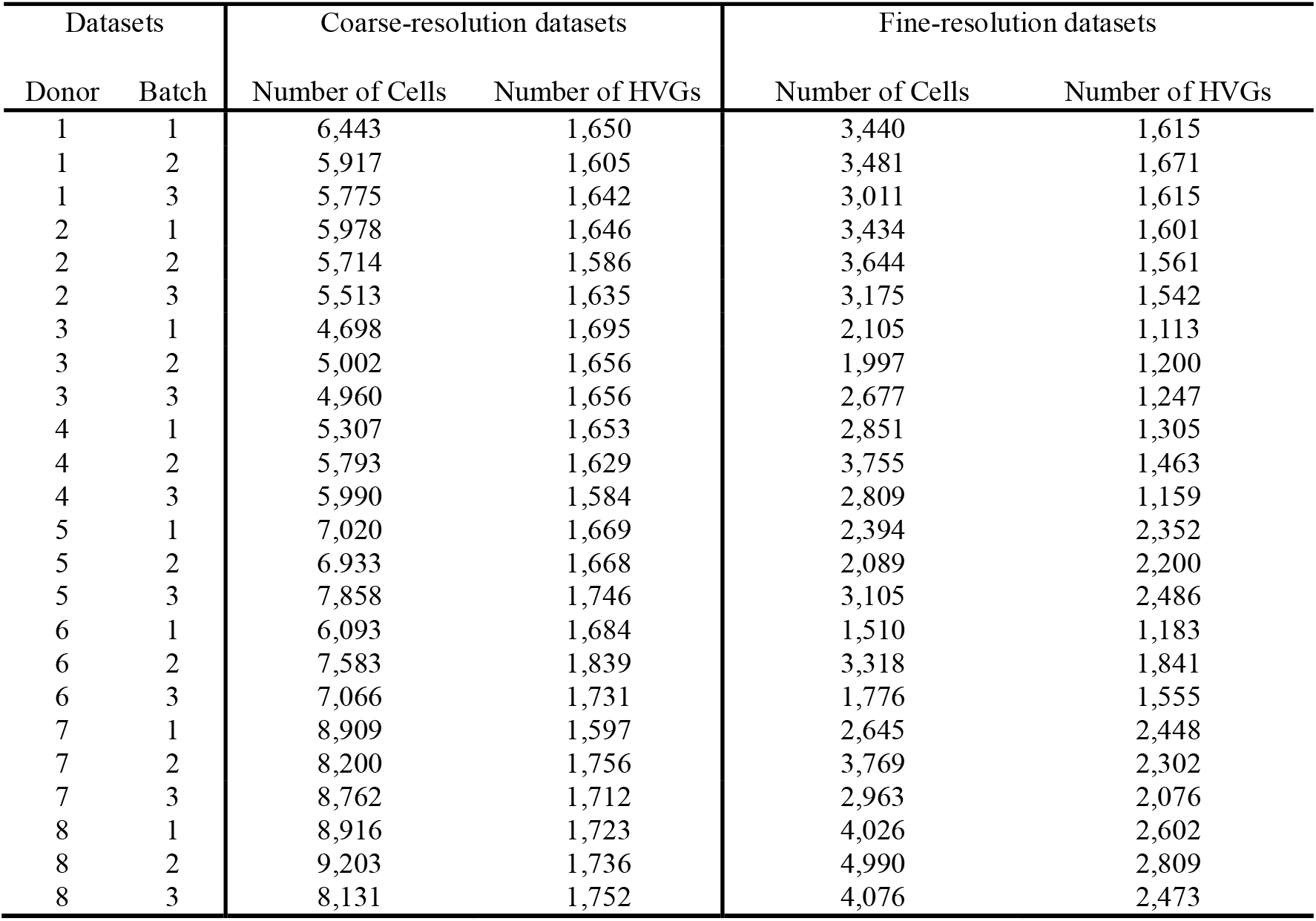
Summary statistics of PBMC datasets.

| Datasets |  | Coarse-resolution datasets |  | Fine-resolution datasets |  |
| --- | --- | --- | --- | --- | --- |
| Donor | Batch | Number of Cells | Number of HVGs | Number of Cells | Number of HVGs |
| 1 | 1 | 6,443 | 1,650 | 3,440 | 1,615 |
| 1 | 2 | 5,917 | 1,605 | 3,481 | 1,671 |
| 1 | 3 | 5,775 | 1,642 | 3,011 | 1,615 |
| 2 | 1 | 5,978 | 1,646 | 3,434 | 1,601 |
| 2 | 2 | 5,714 | 1,586 | 3,644 | 1,561 |
| 2 | 3 | 5,513 | 1,635 | 3,175 | 1,542 |
| 3 | 1 | 4,698 | 1,695 | 2,105 | 1,113 |
| 3 | 2 | 5,002 | 1,656 | 1,997 | 1,200 |
| 3 | 3 | 4,960 | 1,656 | 2,677 | 1,247 |
| 4 | 1 | 5,307 | 1,653 | 2,851 | 1,305 |
| 4 | 2 | 5,793 | 1,629 | 3,755 | 1,463 |
| 4 | 3 | 5,990 | 1,584 | 2,809 | 1,159 |
| 5 | 1 | 7,020 | 1,669 | 2,394 | 2,352 |
| 5 | 2 | 6,933 | 1,668 | 2,089 | 2,200 |
| 5 | 3 | 7,858 | 1,746 | 3,105 | 2,486 |
| 6 | 1 | 6,093 | 1,684 | 1,510 | 1,183 |
| 6 | 2 | 7,583 | 1,839 | 3,318 | 1,841 |
| 6 | 3 | 7,066 | 1,731 | 1,776 | 1,555 |
| 7 | 1 | 8,909 | 1,597 | 2,645 | 2,448 |
| 7 | 2 | 8,200 | 1,756 | 3,769 | 2,302 |
| 7 | 3 | 8,762 | 1,712 | 2,963 | 2,076 |
| 8 | 1 | 8,916 | 1,723 | 4,026 | 2,602 |
| 8 | 2 | 9,203 | 1,736 | 4,990 | 2,809 |
| 8 | 3 | 8,131 | 1,752 | 4,076 | 2,473 |

**Figure S1.**
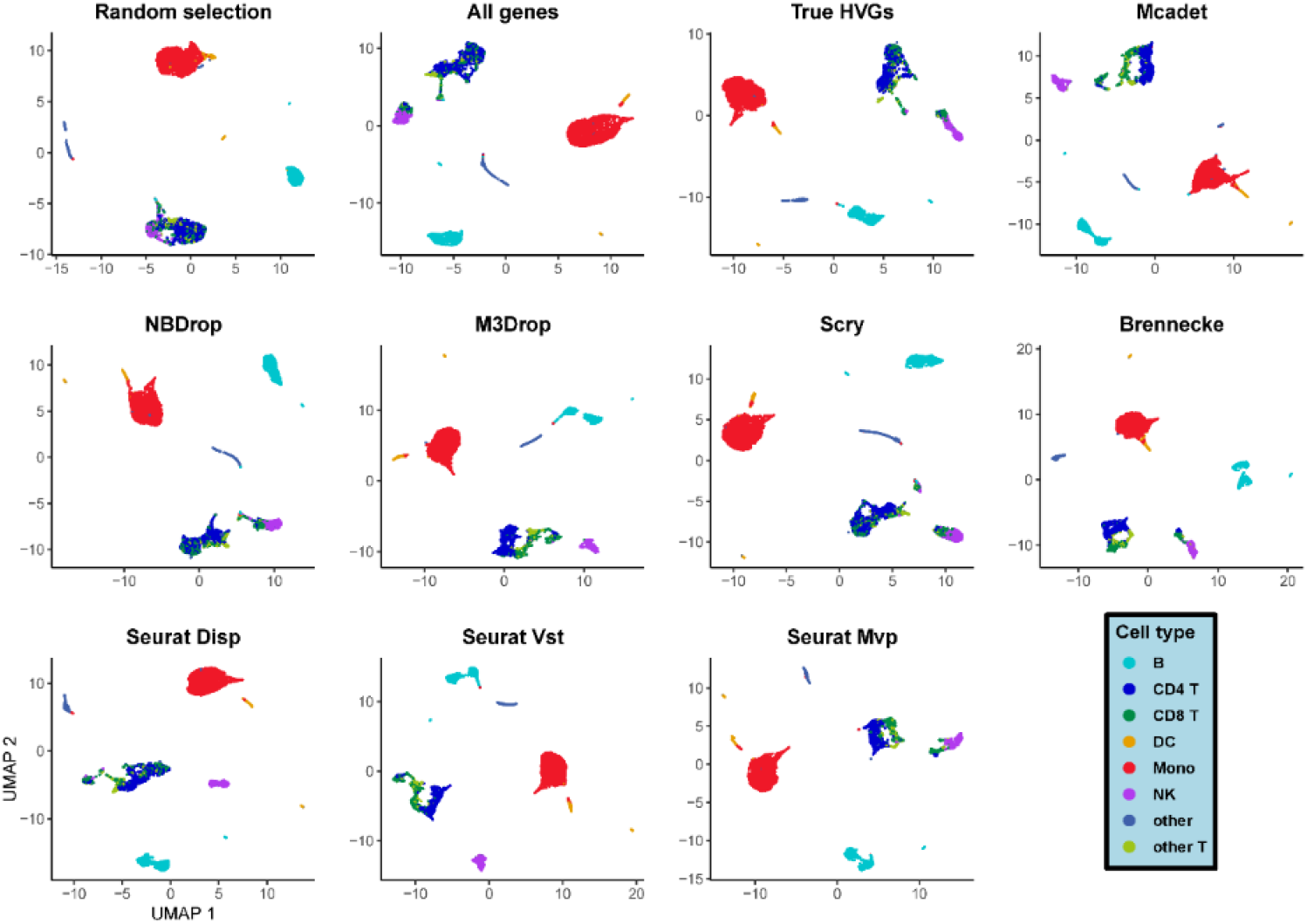
UMAP visualization of a PBMC coarse-resolution scRNA-seq dataset.

**Figure S2.**
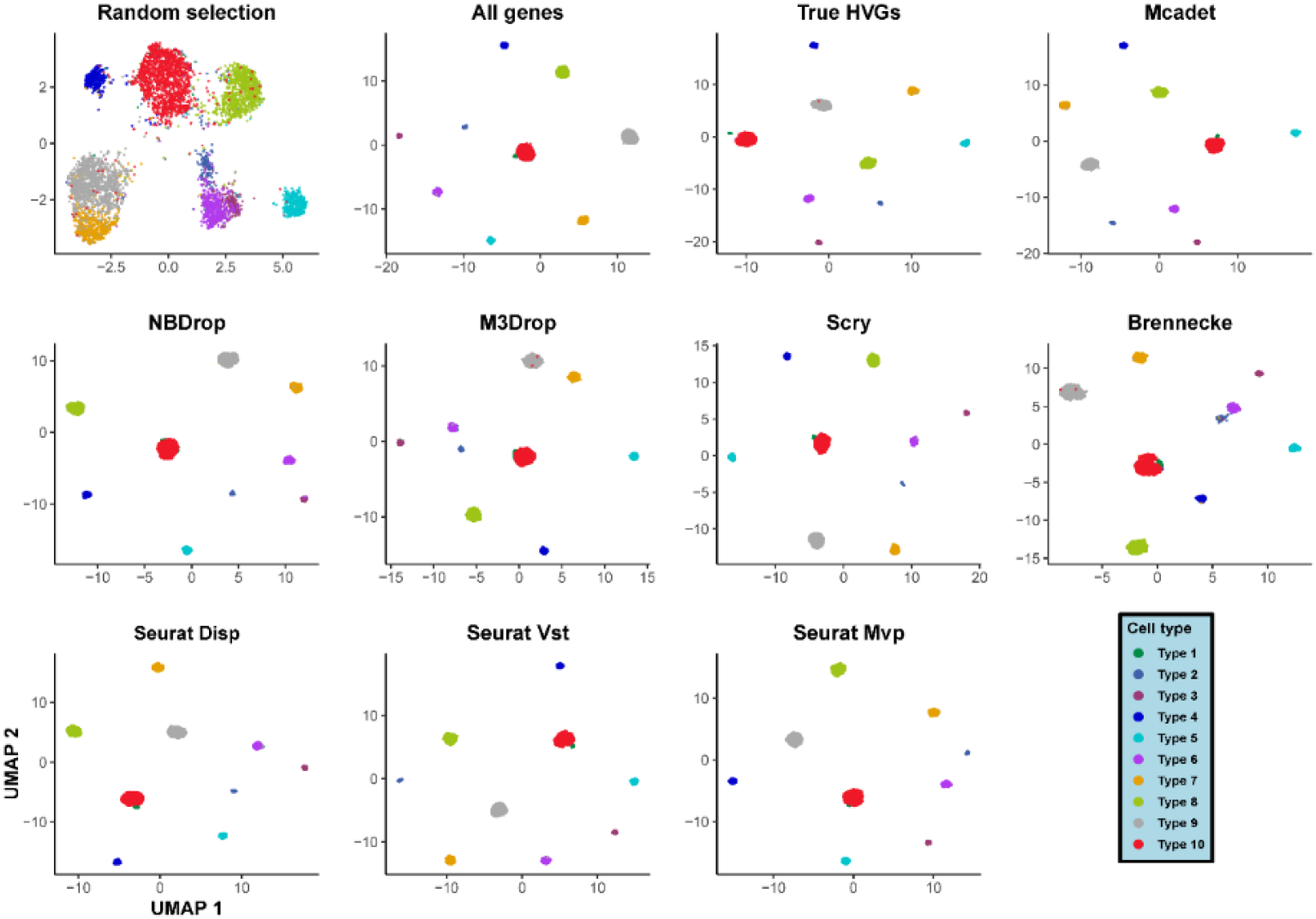
UMAP visualization of a simulated coarse resolution scRNA-seq dataset.

**Figure S3.**
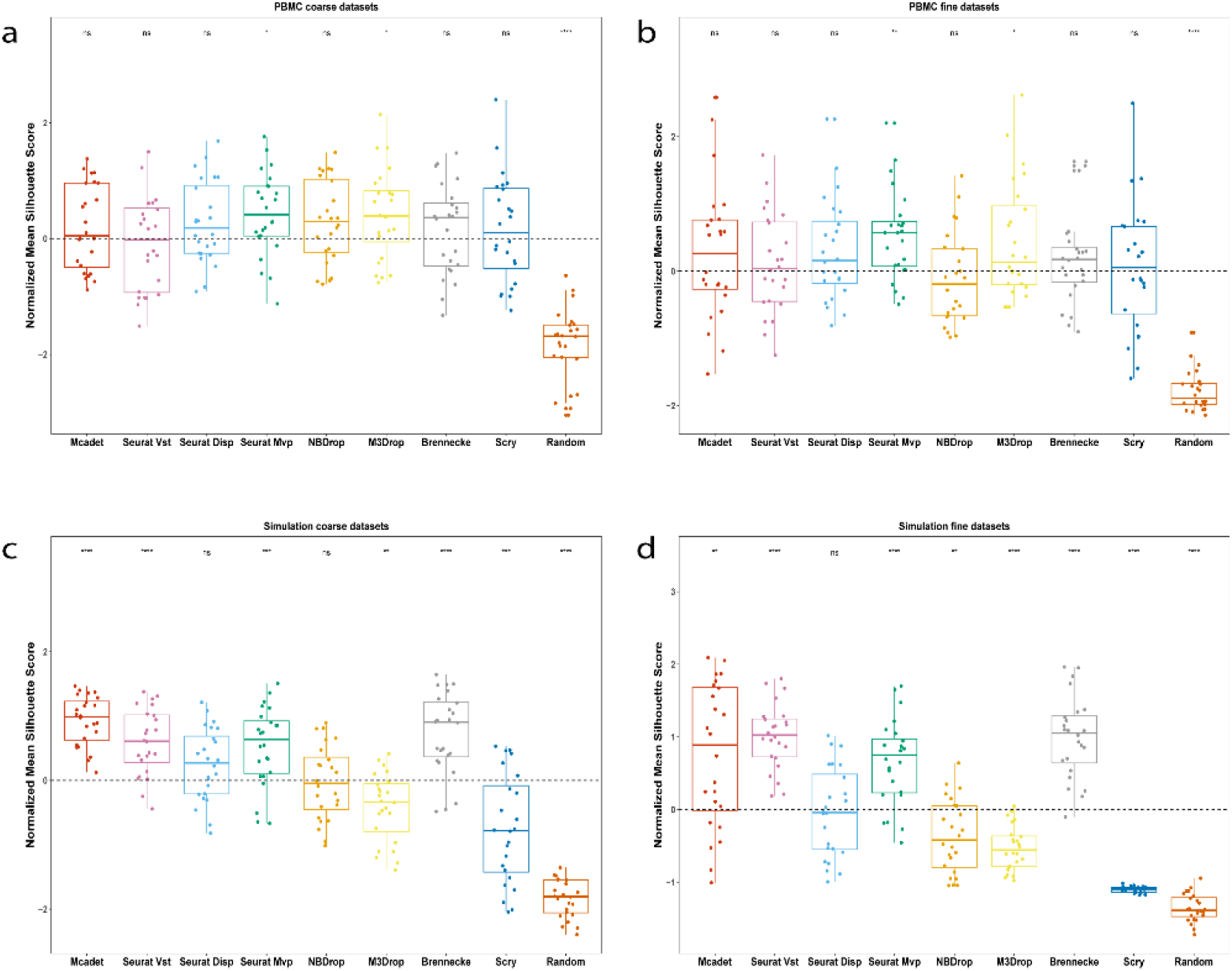
Silhouette score for comparing feature selection performance on PBMC (a-b) and simulated datasets (c-d). Silhouette score is to measure how well the clustering performance with genes selected by different FS methods.

**Figure S4.**
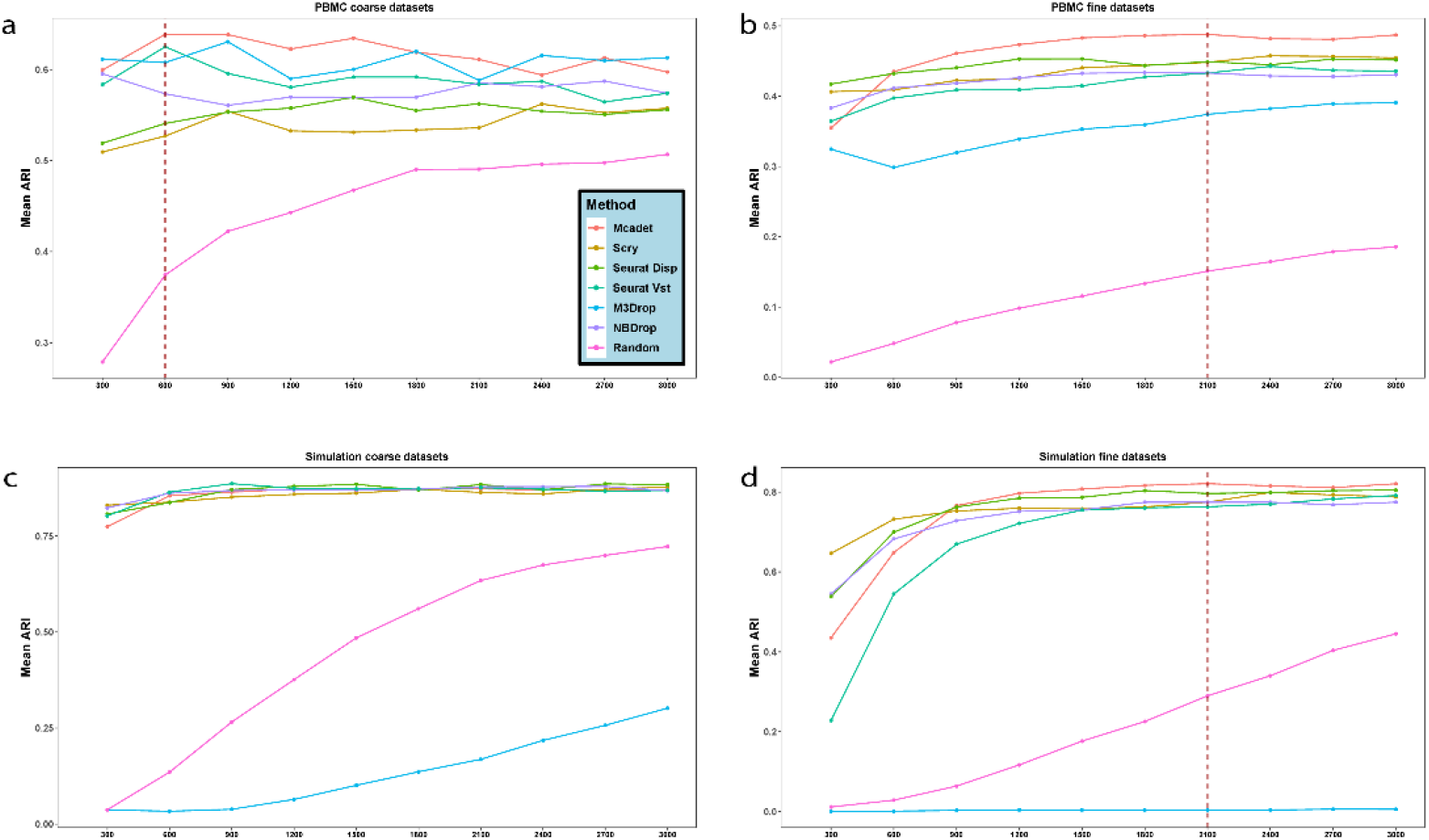
The trend of mean ARI as the number of selected genes increases on PBMC (a-b) and simulated datasets (c-d).

**Figure S5.**
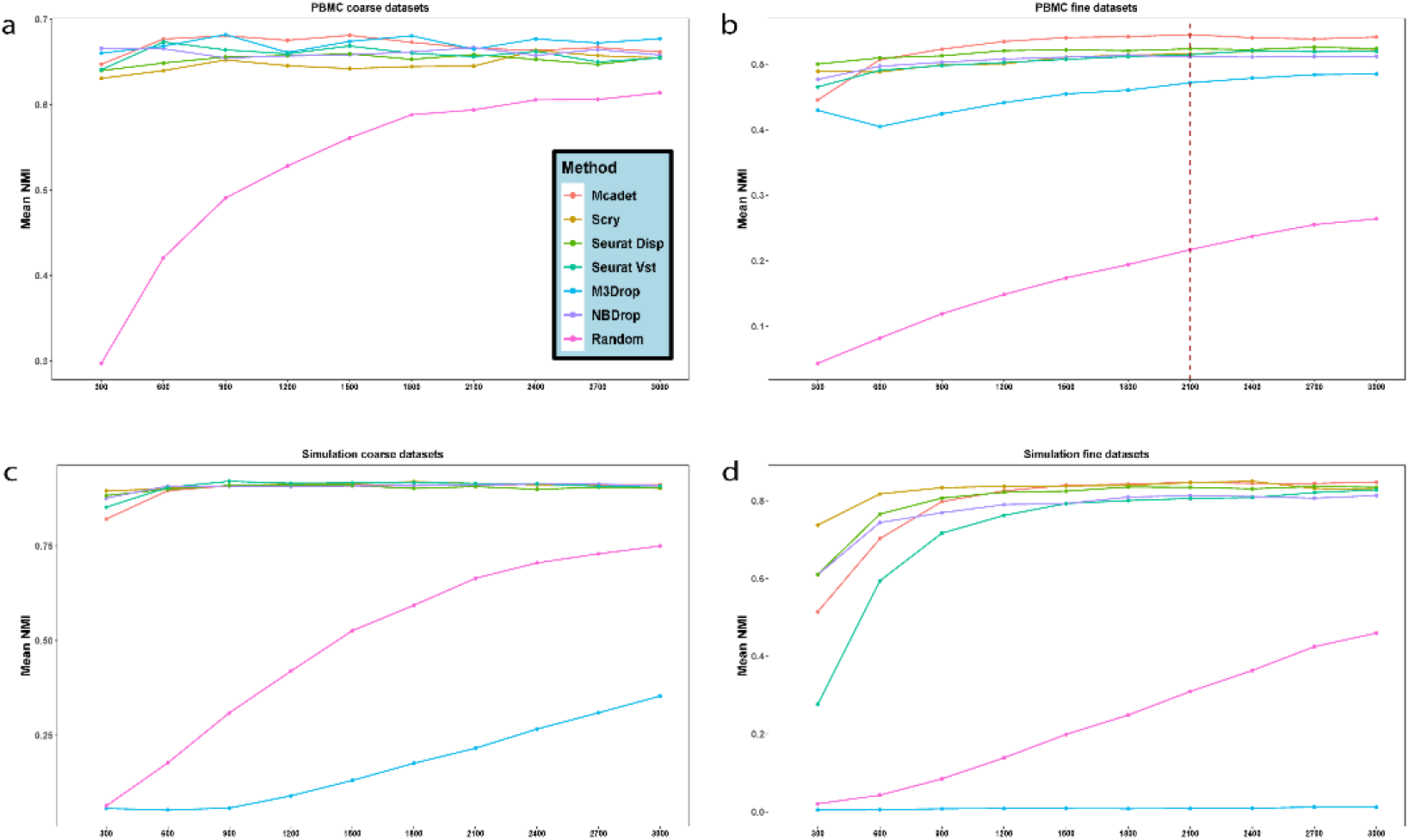
The trend of mean NMI as the number of selected genes increases on PBMC (a-b) and simulated datasets (c-d).

**Figure S6.**
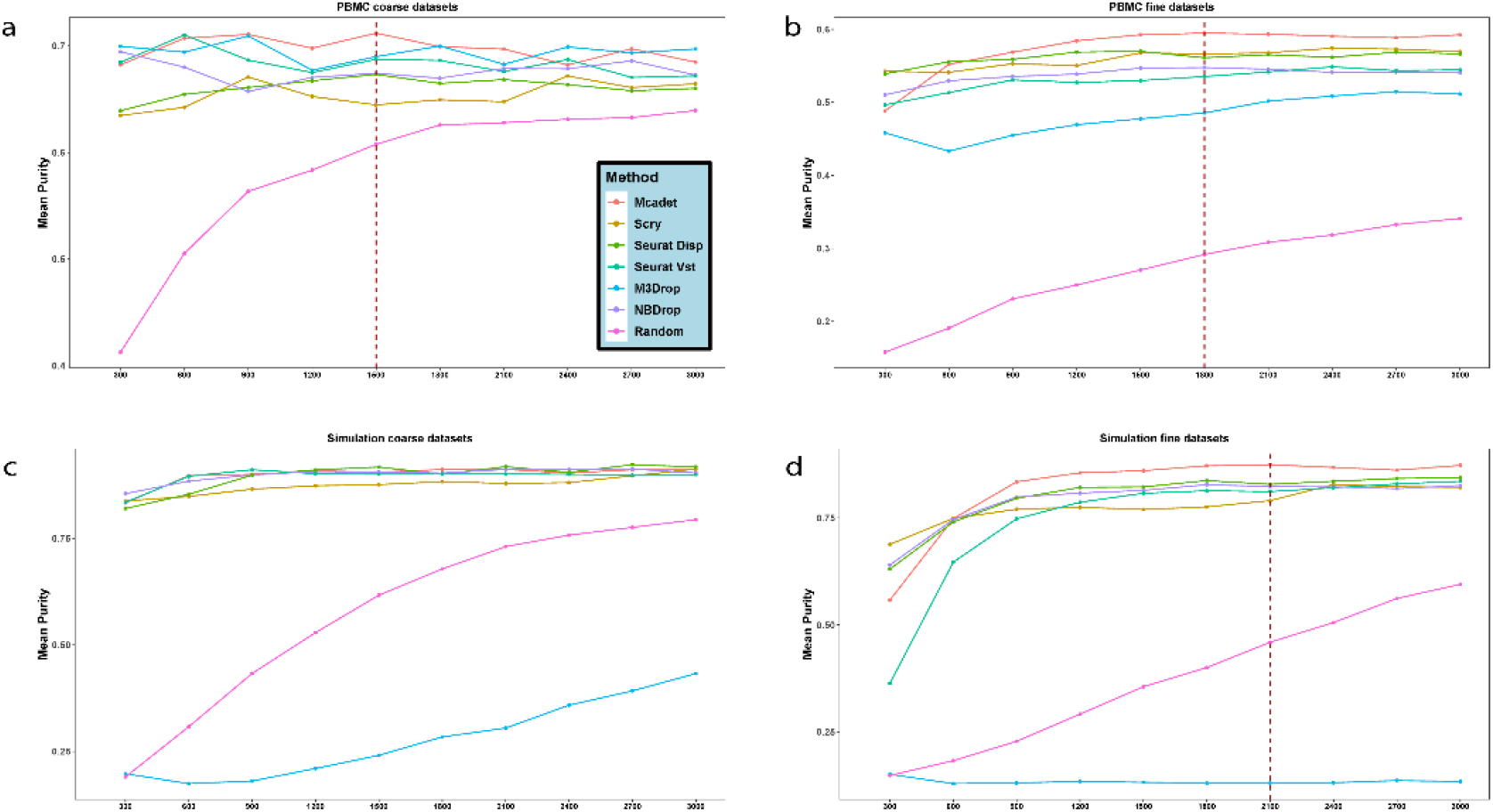
The trend of mean purity as the number of selected genes increases on PBMC (a-b) and simulated datasets (c-d).

**Figure S7.**
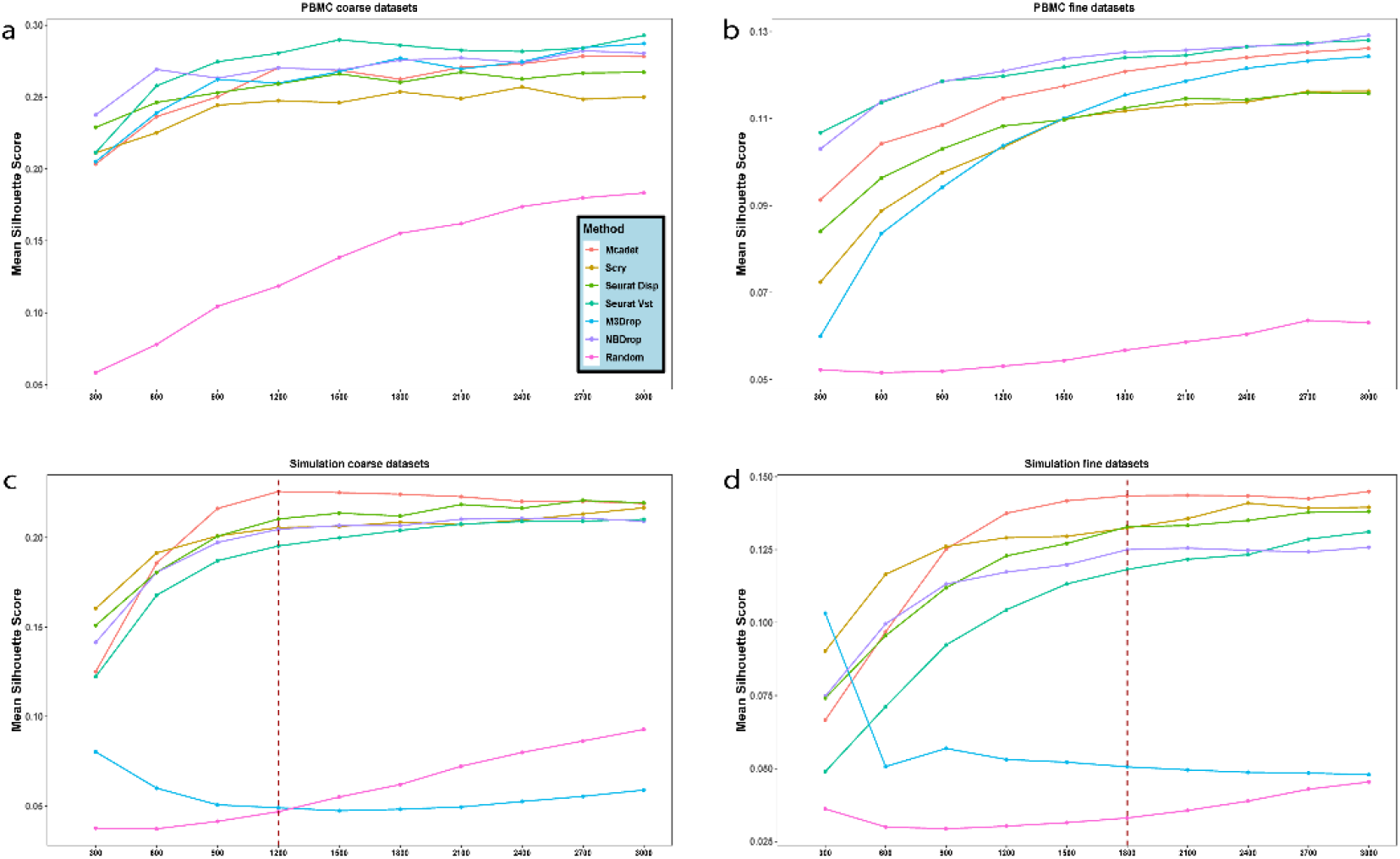
The trend of mean Silhouette score as the number of selected genes increases on PBMC (a-b) and simulated datasets (c-d).

**Figure S8.**
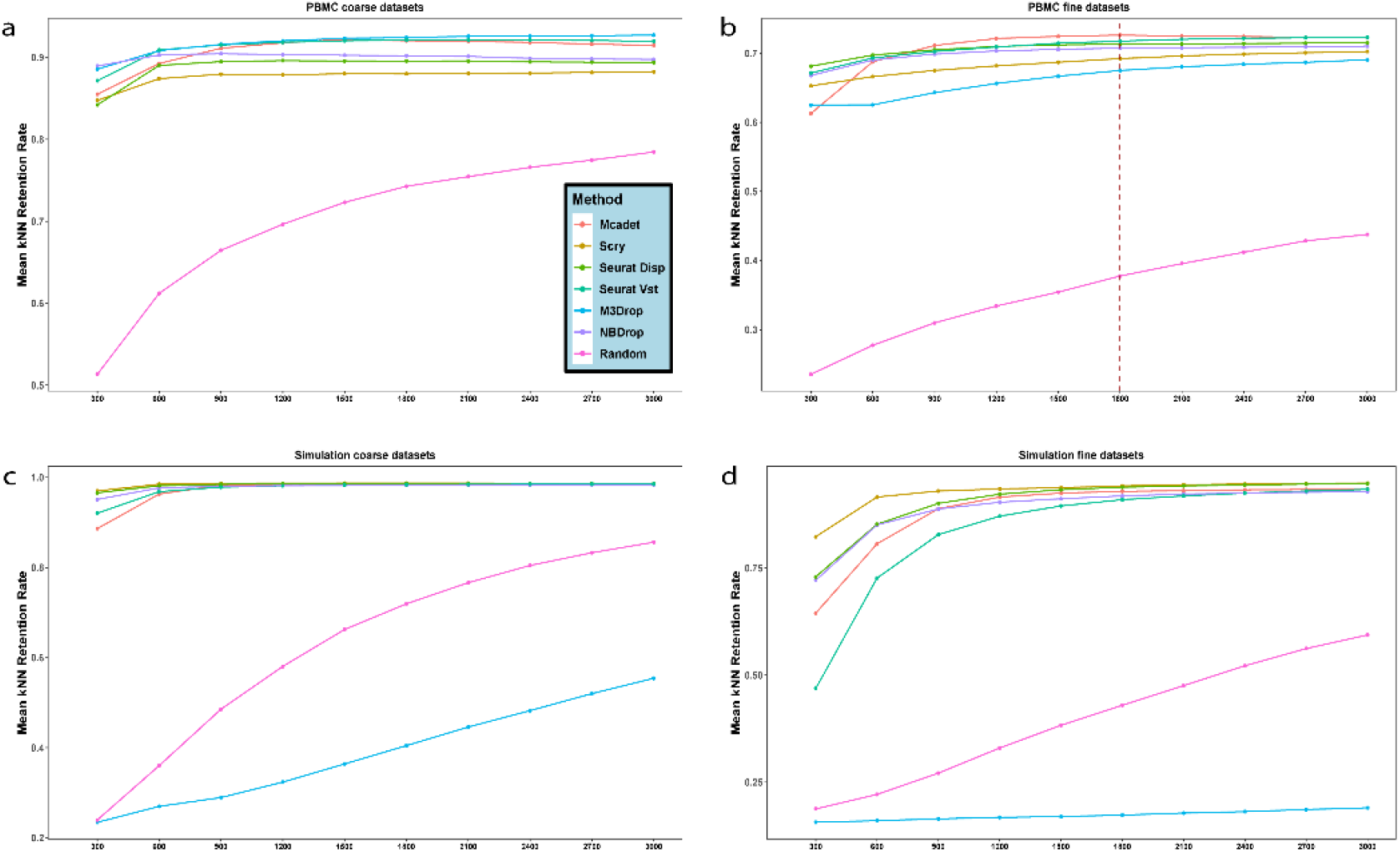
The trend of mean NNgraph retention rate as the number of selected genes increases on PBMC (a-b) and simulated datasets (c-d).

**Figure S9.**
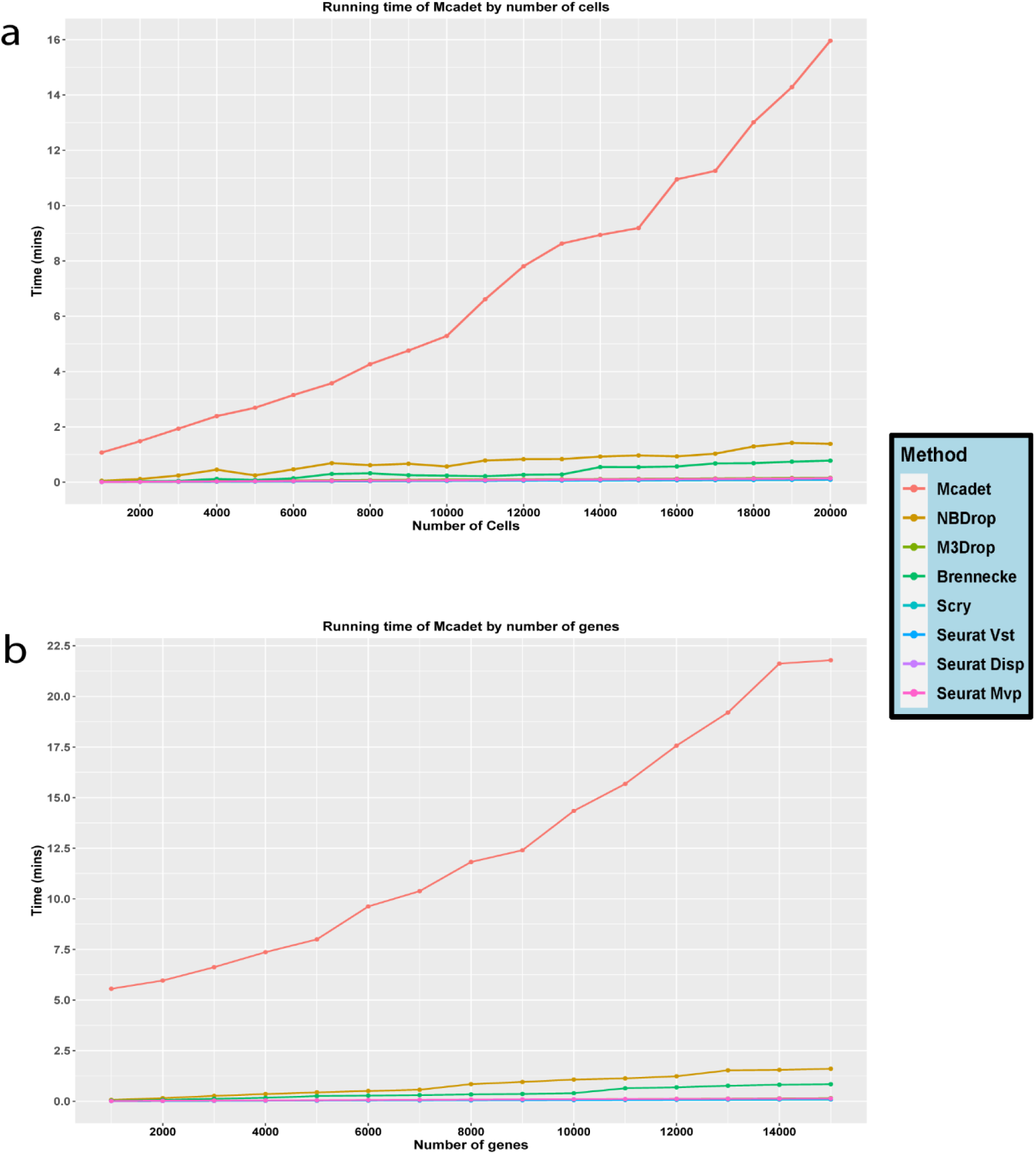
Running time of different FS methods as number of cells (a) and number of genes (b) increase. The experiments were conducted using a simulated dataset on a desktop with an AMD Ryzen 9 7950X 16-Core Processor and 64 GB RAM.

