## Supplementary Methods for "Mcadet: a feature selection method for fine-resolution single-cell RNA-seq data based on multiple correspondence analysis and community detection"

*1. Matrix pre-processing*

For the sake of convenience, we transpose the expression matrix, but this operation does not affect any calculations or interpretations. Let $\boldsymbol{X}_{n\times p}$ be the expression matrix of scRNA-seq data, where $n$ represents the number of cells (rows) and $p$ represents the number of genes (columns). $X_{ij}$ denotes the observed expression of the $j^{th}$ gene in the $i^{th}$ cell.

$$\boldsymbol{X}_{n\times p} = \begin{bmatrix} X_{11} & ... & ... & ... & X_{1p} \\ ... & ... & ... & ... & ... \\ ... & ... & X_{ij} & ... & ... \\ ... & ... & ... & ... & ... \\ X_{n1} & ... & ... & ... & X_{np} \end{bmatrix}_{n\times p} \tag{1}$$

In many scRNA-seq data analyses, it is common to apply a log-normal transformation to the raw count matrix and then perform principal component analysis (PCA) for dimensionality reduction. However, the rationale behind using log counts has limited theoretical justification, and in certain cases, it may obscure meaningful variation. Townes et al. [1] suggested that the zero-inflation resulting from log normalization can be problematic as it artificially amplifies the differences between zero and non-zero values. Booeshaghi et al. [2] demonstrated that for genes with low expression levels, the average log-transformed expression can significantly differ from the expected value $log(\lambda)$, where $\lambda$ is the parameter of the Poisson random variable $X$ being modeled, thus leading to potentially misleading interpretations. Furthermore, Hsu et al. [3] showed that correspondence analysis (CA) remains robust when applied to either count or log-count data, eliminating the need for log-transformation and its associated issues. Therefore, in our method, we retain the raw count matrix for downstream processing.

The fuzzy coding system comprises membership functions and hinge points. By default, the membership function utilized is a triangular function, which is a piecewise linear function. It is important to note that membership functions should be monotonic, as otherwise, the problem would be equivalent to crisp coding, which cannot be converted back to the original data. In our approach, we employ the "doubling" technique in correspondence analysis [4, 5]. This involves separating each variable into two categories. Consequently, the two hinge points serve as endpoints. For the $j^{th}$ gene, $m_{1j}$ (minimum) and $m_{2j}$ (maximum) are used as the hinge points. The membership functions for the "positive" and "negative" doubled variables are simply defined as follows:

$$Z_{ij+} = \frac{X_{ij} - m_{1j}}{m_{2j} - m_{1j}} \in [0, 1], \quad Z_{ij-} = \frac{m_{2j} - X_{ij}}{m_{2j} - m_{1j}} = 1 - Z_{ij+} \in [0, 1] \tag{2}$$

Then after fuzzy coding, the original raw count matrix becomes:

$$\boldsymbol{Z}_{n\times 2p} = \begin{bmatrix} Z_{11+} & Z_{11-} & \cdots & \cdots & \cdots & \cdots & Z_{1p+} & Z_{1p-} \\ \cdots & \cdots & \cdots & \cdots & \cdots & \cdots & \cdots & \cdots \\ \cdots & \cdots & \cdots & Z_{ij+} & Z_{ij-} & \cdots & \cdots & \cdots \\ \cdots & \cdots & \cdots & \cdots & \cdots & \cdots & \cdots & \cdots \\ Z_{n1+} & Z_{n1-} & \cdots & \cdots & \cdots & \cdots & Z_{np+} & Z_{np-} \end{bmatrix}_{n\times 2p} \tag{3}$$

where $Z_{ij+} + Z_{ij-} = 1$, so the total sum is $\sum_{i=1}^{N}\sum_{j=1}^{P} Z_{ij+} + \sum_{i=1}^{N}\sum_{j=1}^{P} Z_{ij-} = NP$.

Fuzzy coding can be regarded as a form of linear transformation that restricts values between 0 and 1, similar to the stabilizing effect of log-normalization. However, one distinctive aspect of fuzzy coding in MCA is that it treats genes with high mean expression and genes with low mean expression similarly, if their presence patterns are comparable. Consequently, both genes contribute approximately equally to the biplot plot. This characteristic sets fuzzy coding MCA apart from other transformations.

*2. MCA decomposition*

Then the correspondence fuzzy coded matrix can be constructed:

$$\boldsymbol{P}_{n\times 2p} = \frac{1}{np}\boldsymbol{Z}_{n\times 2p} = \begin{bmatrix} P_{11+} & P_{11-} & \cdots & \cdots & \cdots & \cdots & P_{1p+} & P_{1p-} \\ \cdots & \cdots & \cdots & \cdots & \cdots & \cdots & \cdots & \cdots \\ \cdots & \cdots & \cdots & P_{ij+} & P_{ij-} & \cdots & \cdots & \cdots \\ \cdots & \cdots & \cdots & \cdots & \cdots & \cdots & \cdots & \cdots \\ P_{n1+} & P_{n1-} & \cdots & \cdots & \cdots & \cdots & P_{np+} & P_{np-} \end{bmatrix}_{n\times 2p} \tag{4}$$

The row mass (margin) and column mass (margin) can be computed:

$$\text{Row mass } r_i = \sum_{j=1}^{p} P_{ij+} + \sum_{j=1}^{p} P_{ij-} = \frac{p}{np} = \frac{1}{n} \tag{5}$$

$$\text{Column mass } c_{j+} = \sum_{i=1}^{n} P_{ij+}, \quad c_{j-} = \sum_{i=1}^{n} P_{ij-} \tag{6}$$

In standard MCA method using indicator (or "pseudo indicator") matrix, the next step is to calculate the standardized Pearson residual matrix with each element

$$r_{ij+/-} = \frac{\text{observed} - \text{expected}}{\sqrt{\text{expected}}} = \frac{P_{ij+/-} - P_{i.}P_{.j+/-}}{\sqrt{P_{i.}P_{.j+/-}}} \tag{7}$$

In matrix notation:

$$\boldsymbol{D_r^{-\frac{1}{2}}(P - rc^T)D_c^{-\frac{1}{2}}} = \boldsymbol{R}_{n\times 2p} = \begin{bmatrix} r_{11+} & r_{11-} & \cdots & \cdots & r_{1p+} & r_{1p-} \\ \cdots & \cdots & \cdots & \cdots & \cdots & \cdots \\ \cdots & \cdots & \cdots & \cdots & \cdots & \cdots \\ \cdots & \cdots & \cdots & \cdots & \cdots & \cdots \\ r_{n1+} & r_{n1-} & \cdots & \cdots & r_{np+} & r_{np-} \end{bmatrix}_{n\times 2p} \tag{8}$$

where $\boldsymbol{r}$ is the vector of row mass, and $\boldsymbol{c}$ is the vector of column mass. $\boldsymbol{D_r}$ is an $n \times n$ diagonal matrix of row mass, and $\boldsymbol{D_r}$ is a $2p \times 2p$ diagonal matrix of column mass. Each element $r_{ij+}$ or $r_{ij-}$ is defined by standardized Pearson residual mentioned above.

Next step is to decompose $\boldsymbol{R}_{n\times 2p}$ by using singular value decomposition (SVD) to find left singular matrix $\boldsymbol{U}$, diagonal matrix of singular values $\boldsymbol{D_\alpha}$ and right singular matrix $\boldsymbol{V}$ such that:

$$\boldsymbol{R}_{n\times 2p} = \boldsymbol{UD_\alpha V^T}, \quad \boldsymbol{U^TU} = \boldsymbol{VV^T} = \boldsymbol{I} \tag{9}$$

In terms of computation method of matrix decomposition, the traditional SVD requires long computation time for large matrix like data matrix in scRNAseq. Therefore, a fast and memory efficient methods of SVD is used, here we employ *irlba* [6] package in R [7].

After SVD of $\boldsymbol{R_{n\times 2p}}$, $K = 60$ (default is 60, one can tune this parameter as well) components are selected as low-dimensional representation. After rearrangements:

$$\boldsymbol{P - rc^T} = \sum_{k=1}^{K} \lambda_k (\boldsymbol{D_r^{\frac{1}{2}} u_k})(\boldsymbol{D_c^{\frac{1}{2}} v_k})^{\boldsymbol{T}} = \sum_{k=1}^{K} \lambda_k (\boldsymbol{D_r D_r^{-\frac{1}{2}} u_k})(\boldsymbol{D_c D_c^{-\frac{1}{2}} v_k})^{\boldsymbol{T}}$$

$$= \boldsymbol{D_r} \left[ \sum_{k=1}^{K} \lambda_k (\boldsymbol{D_r^{-\frac{1}{2}} u_k})(\boldsymbol{D_c^{-\frac{1}{2}} v_k})^{\boldsymbol{T}} \right] \boldsymbol{D_c} \tag{10}$$

Denote standard coordinates of rows and columns as:

$$\boldsymbol{\Phi}_{[\boldsymbol{K}]} = \boldsymbol{D}_{\boldsymbol{r}}^{-\frac{1}{2}}\boldsymbol{U}_{[\boldsymbol{K}]} \text{ and } \boldsymbol{\Gamma}_{[\boldsymbol{K}]} = \boldsymbol{D}_{\boldsymbol{c}}^{-\frac{1}{2}}\boldsymbol{V}_{[\boldsymbol{K}]} \tag{11}$$

Denote principal coordinates of rows and columns as:

$$\boldsymbol{F}_{[\boldsymbol{K}]} = \boldsymbol{D}_{\boldsymbol{r}}^{-\frac{1}{2}}\boldsymbol{U}_{[\boldsymbol{K}]}\boldsymbol{\Lambda}_{[\boldsymbol{K}]} \text{ and } \boldsymbol{G}_{[\boldsymbol{K}]} = \boldsymbol{D}_{\boldsymbol{c}}^{-\frac{1}{2}}\boldsymbol{V}_{[\boldsymbol{K}]}\boldsymbol{\Lambda}_{[\boldsymbol{K}]} \tag{12}$$

The reconstituition formula [8] is given by :

$$\boldsymbol{P} = \boldsymbol{D}_{\boldsymbol{r}}(\boldsymbol{1}_n\boldsymbol{1}_{\boldsymbol{2p}}^{\boldsymbol{T}} + \boldsymbol{\Phi}_{[\boldsymbol{K}]}\boldsymbol{\Lambda}_{[\boldsymbol{K}]}\boldsymbol{\Gamma}_{[\boldsymbol{K}]}^{\boldsymbol{T}})\boldsymbol{D}_{\boldsymbol{c}} \tag{13}$$

Written in column-principal [8] form:

$$\boldsymbol{P} = \boldsymbol{D}_{\boldsymbol{r}}(\boldsymbol{1}_n\boldsymbol{1}_{\boldsymbol{2p}}^{\boldsymbol{T}} + \boldsymbol{\Phi}_{[\boldsymbol{K}]}\boldsymbol{G}_{[\boldsymbol{K}]}^{\boldsymbol{T}})\boldsymbol{D}_{\boldsymbol{c}} \tag{14}$$

An asymmetric map refers to a situation where the row and column points are scaled differently, such as row points being in standard coordinates and column points being in principal coordinates. In contrast, a symmetric map scales the row and column points equally, but it lacks a distance interpretation compared to the asymmetric map [8]. Biplots, which are asymmetric CA maps, combine one set of points in principal coordinates and the other set in standard coordinates. In this study, we utilize a column-principal biplot, which involves using standard row coordinates $\boldsymbol{\Phi}_{[\boldsymbol{K}]} = [\boldsymbol{\phi_1}, \boldsymbol{\phi_2}, \cdots, \boldsymbol{\phi_n}]^T$ and principal column coordinates $\boldsymbol{G}_{[\boldsymbol{K}]} = [\boldsymbol{g_{1+}}, \boldsymbol{g_{1-}}, \cdots, \boldsymbol{g_{p+}}, \boldsymbol{g_{p-}}]^T$ for downstream calculations. Here, $\boldsymbol{\phi_i}$ represents a $K \times 1$ vector of standard row coordinates for cell $i$, and $\boldsymbol{g_{j+/-}}$ represents a $K \times 1$ vector of principal column coordinates for gene $j$. $\boldsymbol{g_{j+}}$ denotes the presence of the gene, while $\boldsymbol{g_{j-}}$ indicates its absence. Each set of two categories for each gene is centered at the origin, that is, $\vec{\boldsymbol{g_{j+}}} + \vec{\boldsymbol{g_{j-}}} = \vec{\boldsymbol{0}}$. Consequently, only the gene category coordinates $\vec{\boldsymbol{g_{j+}}}$ (principal column coordinates), which convey the presence of gene expression relative to the maximum per gene, i.e., $\boldsymbol{G}^*[\boldsymbol{K}] = [\boldsymbol{g1}+, \cdots, \boldsymbol{g_{p+}}]^T$, are retained for downstream analysis.

*3. Community dectection*

Clustering or community detection is performed using the Leiden algorithm, a graph-based clustering algorithm that is faster than the Louvain algorithm. We utilize the R package *leidenAlg* for implementing the Leiden algorithm [9]. Our approach is inspired by previous studies that applied graph-based clustering methods to scRNA-seq data and CyTOF data [10, 11]. These methods involve representing cells as a graph, such as a $k$NN graph, where edges connect cells with similar expression patterns. The objective is to partition the graph into highly interconnected "quasi-cliques" or "communities". Similar to the PhenoGraph method in [11], we construct a KNN graph using the Euclidean distance in the reduced MCA space. We then refine the edge weights between cells based on the shared overlap in their local neighborhoods, measured by the Jaccard similarity. During the construction of the KNN graph, we set the value of $k$ as 1% of the total number of cells in the dataset. To obtain a robust partition, we keep the value of $k$ fixed for building the NNgraph and vary the "resolution" parameter in the find_partition() function of the *leidenAlg* package. We start with a resolution of 0.5 and increment it by 0.1, running the algorithm 10 times by default.

*4. Calculating and ranking the Euclidean distance*

Having established the cell communities (or clusters), assume there are $d$ clusters, a set of cells $\Psi \subseteq \{C_1, \cdots, C_d\}$ can be defined, we can calculate the coordinates of each cluster centroid by taking the average coordinates of the cells in a cluster:

$$\boldsymbol{T_\Psi} = \frac{1}{|\Psi|} \cdot \sum_{i \in \Psi} \boldsymbol{\phi_i} \tag{15}$$

where $\boldsymbol{T_\Psi}$ is a $K \times 1$ vector, the coordinates of cell community $\Psi$, $|\Psi|$ is the number of cells in community $\Psi$, and $\boldsymbol{\phi_i}$ is the coordinates of cell $i$ in this low-dimensional space we defined above.

Once we have obtained the coordinates of each cluster centroid in the low-dimensional MCA space, we can calculate the Euclidean distances between each gene $(+)$ and each cell centroid. In this low-dimensional MCA biplot representation, the distance between a column (representing a gene $+$) and a row (representing a cell) indicates the specificity of that gene to the corresponding cell. A shorter distance suggests a higher specificity. Therefore, to assess the specificity of a gene to a particular cell community, we calculate the Euclidean distance between each gene and each community centroid.

$$d_{(j,\Psi)} = \sqrt{(\boldsymbol{g_{j+}} - \boldsymbol{T_\Psi})^{\boldsymbol{T}}(\boldsymbol{g_{j+}} - \boldsymbol{T_\Psi})} \tag{16}$$

where $d_{(j,\Psi)}$ is the Euclidean distance between gene $j(+)$ and cell community centroid $\Psi$, and $\boldsymbol{g_{j+}}$ is the coordinates of gene $j$ in this low-dimensional space we defined above. $\boldsymbol{T_\Psi}$ is the coordinate of cell community $\Psi$.

Next, based on the Euclidean distances we have obtained, we can rank them by each community:

$$\{rd_{(1,\Psi)}, \cdots, rd_{(p,\Psi)}\} = rank\{d_{(1,\Psi)}, \cdots, d_{(p,\Psi)}\}, \quad \Psi \subseteq \{C_1, \cdots, C_d\} \tag{17}$$

where $rd_{(j,\Psi)}$ is the rank of gene $j$ in the Euclidean distances between all genes to cell community $\Psi$'s centroid. We utilize ranks instead of the actual distances because the Euclidean distances can vary significantly in magnitude due to the cells being distributed across an irregular and high-dimensional space. The distances do not follow a normal distribution, making parametric models such as ANOVA less effective. This limitation is also present in the FEAST method, which employs F-statistics to assess feature significance based on mean gene expression [12]. However, scRNA-seq data often deviate from normal distributions, even after log normalization, primarily due to the high fraction of zero counts [1, 2]. Nonetheless, the principle that "the closer a column is to a row, the more specific it is to it" can be applied in any high-dimensional space, regardless of the "curse of dimensionality", and is not confined to any specific distribution. Thus, the rank of distances for genes within each cluster holds meaningful information. Consequently, we can generate a rank list $\Omega_j$ for each gene:

$$\Omega_j = \{rd_{(j,C_1)}, \cdots, rd_{(j,\Psi)}, \cdots, rd_{(j,C_d)}\} \tag{18}$$

The maximum and minimum order statistics of $\Omega_j$, denoted as $max(\Omega_j)$ and $min(\Omega_j)$, provide insights into gene variability. For instance, consider gene $m_1$ with $min(\Omega_{m_1})$ equal to 5, indicating that it ranks significantly higher in a particular cell community compared to other genes. This suggests that gene $m_1$ is highly expressed in this specific cell community. Conversely, its $max(\Omega_{m_1})$ is $10,000$, indicating that gene $m_1$ is not prominently expressed in another population of cells compared to other genes. The log ratio between $max(\Omega_{m_1})$ and $min(\Omega_{m_1})$, i.e., $log(10000/5) = 7.6$, is relatively large. Thus, gene $m_1$ can be considered a candidate driver gene in our feature selection task. In contrast, let's consider gene $m_2$, where $min(\Omega_{m_2})$ is $12,000$ and $max(\Omega_{m_2})$ is $14,000$. These values indicate that gene $m_2$ is distanced from any cell clusters compared to other genes. The log ratio between $max(\Omega_{m_2})$ and $min(\Omega_{m_2})$ is only 0.15, suggesting that gene $m_2$ is unlikely to be selected as a feature. Similarly, a housekeeping gene $m_3$, which is expressed in the majority of cell types, would be close to any of the cell clusters. In this case, we have $min(\Omega_{m_3}) = 200$ and $max(\Omega_{m_3}) = 800$, resulting in a log ratio of 1.4, which is also small compared to $m_1$. Therefore, gene $m_3$ would not be chosen as a feature in our method.

Hence we define the metric as $logR$:

$$logR_j = log\left(\frac{max(\Omega_j)}{min(\Omega_j)}\right) \tag{19}$$

Then we rank $logR_j$ from high to low, and get top $\gamma$ genes specified by users who know how many genes they want:

$$\Gamma = \{g_j : |\forall j \in \{1, \cdots, p\} : \underset{j}{rank}(logR_j, \text{descending}) \leq \gamma\} \tag{20}$$

where $\Gamma$ is the final gene set selected. $\gamma$ is the parameter can be tuned by users who have domain knowledge. Descending stands for ranking from high to low.

*5. Statistical testing*

The approach described above requires users to specify the number of informative genes to be selected, which relies on domain knowledge and may vary across different samples. However, since the number of genes detected can differ among samples, a more data-driven method is required.

Suppose after Leiden community detection, there are $d$ clusters detected, and suppose there are $p$ detected genes, and for gene $j$, suppose the rank pattern is: $\{X_{1j}, \cdots X_{ij}, \cdots X_{dj}\}$,

$X_{ij}$ is an integer in 1 to $p$, indicating the Euclidean distance rank for gene $j$ in cluster $i$. Our assumption is that for each gene the random variable of rank pattern $X_j$ is i.i.d., and follows uniform distribution (discrete).

$$P(X_j = k) = \frac{1}{p}, \quad 1 \le k \le p \tag{21}$$

Suppose $X_{(1)} < X_{(2)} < \cdots < X_{(d)}$ are the order statistic of random variable $X$. We can derive the joint distribution of minimum $X_{(1)}$ and maximum $X_{(d)}$ as:

$$P(X_{(1)} = k_1, X_{(d)} = k_2) = \left(\frac{k_2 - k_1 - 1}{p}\right)^d + \left(\frac{k_2 - k_1 + 1}{p}\right)^d - 2\left(\frac{k_2 - k_1}{p}\right)^d \tag{22}$$

Proof of (22):
According to the order statistic and (21), we know that:

$$P(X_{(d)} \le k) = \left(\frac{k}{p}\right)^d$$

then,

$$P(X_{(d)} = k) = P(X_{(d)} \le k) - P(X_{(d)} \le k - 1) = \left(\frac{k}{p}\right)^d - \left(\frac{k-1}{p}\right)^d$$

and

$$P(X_{(1)} \ge k_1, X_{(d)} \le k_2) = P(k_1 \le X_{(1)} \le k_2) \cdot P(k_1 \le X_{(2)} \le k_2) \cdots P(k_1 \le X_{(d)} \le k_2)$$

$$= \left(\frac{k_2 - k_1}{p}\right)^d$$

Based on the above formula, we can derive the joint distribution of $X_{(1)}$ and $X_{(d)}$:

$$P(X_{(d)} \le k_2) = P(X_{(1)} \le k_1, X_{(d)} \le k_2) + P(X_{(1)} \ge k_1, X_{(d)} \le k_2)$$

$$\Longrightarrow P(X_{(1)} \le k_1, X_{(d)} \le k_2) = P(X_{(d)} \le k_2) - P(X_{(1)} \ge k_1, X_{(d)} \le k_2)$$
$$= \left(\frac{k_2}{p}\right)^d - \left(\frac{k_2 - k_1}{p}\right)^d$$

then,

$$P(X_{(1)} \le k_1, X_{(d)} = k_2) = P(X_{(1)} \le k_1, X_{(d)} \le k_2) - P(X_{(1)} \le k_1, X_{(d)} \le k_2 - 1)$$

$$= \left(\frac{k_2}{p}\right)^d - \left(\frac{k_2-k_1}{p}\right)^d - \left(\frac{k_2-1}{p}\right)^d + \left(\frac{k_2-k_1-1}{p}\right)^d$$

and,

$$P(X_{(1)} = k_1, X_{(d)} = k_2) = P(X_{(1)} \leq k_1, X_{(d)} = k_2) - P(X_{(1)} \leq k_1 - 1, X_{(d)} = k_2)$$

$$= \left(\frac{k_2-k_1-1}{p}\right)^d + \left(\frac{k_2-k_1+1}{p}\right)^d - 2\left(\frac{k_2-k_1}{p}\right)^d$$

Therefore, (22) has proved.

Let $R = \frac{X_{(d)}}{X_{(1)}}$, $U = X_{(1)} \implies X_{(1)} = U, X_{(d)} = UR$. Using bivariate transformation to get the joint distribution of $R$ and $U$:

$$P(R = r, U = u) = P(X_{(1)} = U, X_{(d)} = UR)$$

$$= \left(\frac{ur-u-1}{p}\right)^d + \left(\frac{ur-u+1}{p}\right)^d - 2\left(\frac{ur-u}{p}\right)^d$$

Then we can get the marginal distribution of $R$: $P(R = r) =$

$$\sum_{u:\{ur\in Z^+, ur\leq p\}} P(R = r, U = u) = \sum_{u:\{ur\in Z^+, ur\leq p\}} \left[\left(\frac{ur-u-1}{p}\right)^d + \left(\frac{ur-u+1}{p}\right)^d - 2\left(\frac{ur-u}{p}\right)^d\right]$$

Let $V = logR = log\frac{X_{(d)}}{X_{(1)}}$ as the log ratio between the maximum and the minimum rank. The distribution of $V$ can be derived,

$$P(V = v) = \sum_{u\in\boldsymbol{U}} \left[\left(\frac{ue^v-u-1}{p}\right)^d + \left(\frac{ue^v-u+1}{p}\right)^d - 2\left(\frac{ue^v-u}{p}\right)^d\right] \quad (23)$$

where $\boldsymbol{U} = \{u : ue^v \in Z^+, ue^v \leq p\}$

This implies that if our assumption (null hypothesis) holds, stating that ranks are chosen with equal probability for each position, the random variable $V = \log\left(\frac{X_{(d)}}{X_{(1)}}\right)$ will follow the distribution described in equation (23) as $Rdis(p, d)$, with two parameters $p$ and $d$. Let $Q(V = v) = P(V \leq v)$ be the cumulative distribution function defined in equation (23). For a specific gene, if its observed $V$ value is $\tilde{v}$ and $Q(\tilde{v}) > 1 - \alpha$, where $\alpha$ is the type I error typically set at 0.05, we can reject the null hypothesis. This suggests that the observed maximum or minimum rank is not randomly selected from the range of 1 to $p$. Instead, it indicates that the gene is more likely to have a higher probability of being assigned the maximum rank among larger numbers or the minimum rank among smaller numbers. This observation implies that the gene may carry informative characteristics.

In real datasets, it is important to acknowledge that the variables $X_j$ are not independent. This is because the presence of one cluster can influence another cluster. While this assumption is strong, it provides a simple model to work with. Furthermore, if we reject the null hypothesis $H_0$, which is based on the assumption of independent and identically distributed (i.i.d.) variables $X_j$, it suggests that the true distribution of $X_j$ under gene $j$ does not follow equation (21), and therefore, the clusters may not be independent of each other. Additionally, when dealing with a large number of variables $p$ (which is typically greater than 10,000 in real datasets), there will be a vast number of possible observed $v$ values, given by $v = \log\left(\frac{X_{(d)}}{X_{(1)}}\right)$. For instance, if $p = 15,000$, there would be approximately 112,507,500 (about 0.1 billion) possible observations. After removing duplicates, we are left with approximately 68,394,316 (68 million) unique $v$ values. Considering this, if we obtain an observation $\tilde{v}$, calculating its p-value using the exact probability mass function from equation (23) would be time-consuming, as it requires summing up probabilities one by one due to the discrete nature of the distribution. To address this, we employ Monte Carlo simulation to estimate p-values. In each simulation iteration, we randomly draw a rank from 1 to $p$ and compute $v$ as $\log\left(\frac{X_{(d)}}{X_{(1)}}\right)$. This process is repeated $T$ times, where $T$ is typically set to a large number such as 20,000 or 30,000, in order to generate a simulated distribution. Consequently, p-values can be obtained by comparing the observed $\tilde{v}$ to the simulated distribution.

Since each time we input a different resolution parameter for Leiden community detection, we may obtain a different number of clusters $d$ from Leiden community detection. When the Leiden algorithm is run $t$ times, for time $t$ we detect $d_t$ clusters, then, under the null hypothesis, the overall log ratio between maximum rank and minimum rank will follow the distribution:

$$\text{Overall log ratio} \sim Rdis(p, d_1) + Rdis(p, d_2) + \cdots + Rdis(p, d_t) \tag{24}$$

Observed log ratios can be calculated by summing all observed data, that is, overall $\tilde{v} = \tilde{v}_1 + \tilde{v}_2 + \cdots + \tilde{v}_t$. The upper-tailed p-value of each gene is then determined by using the Monte Carlo simulation described above.

*6. Multiple-testing issue*

To address the issue of multiple testing, we will utilize the Benjamini-Hochberg (BH) procedure [13] to control the false discovery rate. It is important to note that the BH procedure assumes that p-value statistics follow a uniform distribution under the null hypotheses. However, in the case of discrete distributions for the test statistics, this assumption is violated. Our test statistic also follows a discrete distribution. To overcome this limitation, Joshua and Edsel [14] proposed a stochastic process framework that enables p-value statistics to satisfy the condition of uniformity by introducing a uniform variate for test statistics with discrete distributions. However, in our testing scenario, this correction for p-values has minimal impact. Despite the discrete nature of our test statistic's distribution, the large number of possible values for the test statistics mentioned earlier results in an approximately continuous distribution. Moreover, scRNA-seq datasets typically contain numerous genes with upper-tailed p-values close to 1, which violates the assumption of uniformity. Therefore, we remove genes with p-values greater than 0.9. Consequently, the remaining p-values under the null hypothesis will be approximately uniformly distributed, allowing us to apply the BH procedure.

*7. Default tuning parameters of Mcadet function*

The Mcadet function implements a computational pipeline for scRNA-seq data analysis. It contains six key parameters that can be specified by the user:

1). n.comp: The number of PCs to use for dimensionality reduction in the initial MCA. The default is set to 60 PCs.

2). run: The number of iterations need to be ran to get a more robust result. The default is 10 iterations.

3). start_resolution: The starting resolution parameter for the Leiden algorithm in each iteration. The default is 0.5, incremented by 0.1 in each subsequent iteration.

4). cell_percent: The percentage of cells a gene must be expressed in for it to be included in the analysis. Genes expressed in fewer cells are excluded as non-informative. The

default threshold is 0.5% of cells.
5). MC_iter: The number of Monte-Carlo iterations for statistical testing of cluster-specific gene expression. The default is 50,000 iterations.
6). fdr: The false discovery rate used as threshold for identifying significant genes. The default FDR is set to 0.15.

In summary, Mcadet allows customization of key parameters controlling dimensionality reduction, clustering, and differential expression analysis within the computational pipeline. The defaults are empirically calibrated for robust performance across diverse scRNA-seq datasets.
